# Mapping of long-term impact of conventional and organic soil management on resident and active fractions of rhizosphere communities of barley

**DOI:** 10.1101/546192

**Authors:** Paula Harkes, Afnan K.A. Suleiman, Sven J.J. van den Elsen, Janjo J. de Haan, Martijn Holterman, Eiko E. Kuramae, Johannes Helder

## Abstract

Soil biota plays an essential role in ecosystem services such as carbon fixation, nitrogen and phosphorous cycling, and disease suppressiveness. Conventional soil management with large inputs of mineral fertilizers and pesticides have a significant impact on primary decomposer communities (bacteria and fungi), as well as on protists and metazoa, representatives of the next trophic level. Organic soil management is thought to contribute to a more diverse and stable soil food web. However, information to pinpoint this supposed beneficial effect is sparse and fragmented. Keeping in mind that a substantial fraction of the soil biota is dormant, we set out to map both the resident and the active the bacterial, fungal, protozoan and metazoan communities under various soil management regimes in two distinct soil types with barley as main crop. For all four organismal groups, the contrast between resident (rDNA-based) and active (rRNA-based) was the most important explanatory variable explaining 22%, 14%, 21% and 25% of the variance among bacterial, fungal, protozoan, and metazoan communities. Less prominent were the effects of soil management and soil type, however significant as well for all four organismal groups. LEfSe was used to identify indicator taxa for both the contrasts between resident and active communities, and the effects of soil management. Our results suggest that - next to DNA-based community characterisation - mapping of the active microbial community could provide essential insights in the effects of variables such as crop and soil management on the soil living community.

## Introduction

Over the last century, rising food demands have been met with an expansion of the cultivated areas as well as with an intensification of agricultural practices. This intensification process is associated with increased inputs such as increased fertilization, intense pesticide applications, frequent tilling, and improved water management (Tilman et al. 2002) However, the negative environmental implications of this so-called green revolution are widely recognised, and include eutrophication, increased greenhouse gas emission, soil erosion and biodiversity loss (Foley et al. 2005, Tilman et al. 2001). Aboveground biodiversity loss is intensively studied, however our knowledge on the impact of agricultural intensification on belowground biodiversity is still fragmented (Hole et al. 2005). It is widely acknowledged that the nutritional needs of an ever-increasing world population can only be addressed structurally by developments towards more sustainable crop production. As one of the consequences, more emphasis is put on understanding and unravelling the biological functioning of soil, being one of the most essential and complex components of our agroecosystems (Tilman et al. 2011).

Soil harbours a quarter of the world’s biodiversity and is considered to be one of the most complex ecosystems on earth (Bardgett 2005, FAO 2015). In terms of numbers of individual organism per unit of volume, bacteria are by far the most abundant group of soil inhabitants followed by fungi, algae, protists and nematodes (Fierer et al. 2007). Soil biota plays a role in many essential soil functions such as nutrient cycling, carbon and water retention, soil texture formation and the interaction with the plant community (Henneron et al. 2015). The most intensive interactions between microbes and plants take place at the rhizosphere, which is the interface between plant roots and their surrounding soil. In the rhizosphere, the plant is able to select and boost a subset of microorganisms by the release of rhizodeposits - a broad range of carbon-containing substances (e.g. root cells, mucilage, volatiles and exudates). As carbon is often the limiting factor for growth of microorganisms, rhizodeposits are thought to be an important driver of shaping soil microbial community structure (Birkett et al. 2001, Paterson et al. 2007). Microorganisms in return promote mineral turnover and/or may play a role in the protection of the plants against pathogens (Berendsen et al. 2012, Lugtenberg & Kamilova 2009). The composition of the rhizobiome, the subset of the soil biota present in the rhizosphere, is co-determined by plant species and age (Chaparro et al. 2014, Grayston et al. 1998, Inceoglu et al. 2011, Kuske et al. 2002, Schlemper et al. 2017). With the advent of affordable high throughput DNA sequencing techniques, the impact of plants on the identity and density of rhizosphere inhabitants can be mapped. Insight in this process could help to design soil management measures promoting a rhizobiome that would optimally support plant growth and improve crop yield (Knief 2014).

Just like in many other habitats, most soil inhabitants have to cope with unpredictable food availability (Morita & Morita 1997). In order to survive periods of scarcity, microorganisms can reversibly reduce their metabolic activity over an extended period of time (Blagodatskaya & Kuzyakov 2013). Such a condition is referred to as a state of dormancy (Stevenson 1978). In bulk soil, typically 80% of the cells and 50% of the operational taxonomical units (OTU’s) are dormant. This so-called “microbial seedbank” (Lennon & Jones 2011) is alert in the sense that it can detect and respond to environmental stimuli (e.g. organic substrates) that are associated with favourable growing conditions (De Nobili et al. 2001). Plant roots produce and release a broad spectrum of environmental stimuli and, as such, the rhizosphere is a hotspot of microbial activity (Hinsinger et al. 2009, Reinhold-Hurek et al. 2015).

In this respect, it is relevant to discriminate between the resident and the active microbial community. The resident community refers here to all organisms present in a certain spatial unit of soil, whereas the active community comprises the fraction of the resident community that is metabolically active. The RNA is considered a representation of the active microbial community, while the DNA characterises the total microbial community (De Vrieze et al. 2016, Ofek et al. 2014). Hence, combined profiling of community DNA and RNA will provide insight in both aspects of local microbial assemblies. More specifically, such a characterisation will provide information about microbial fractions, which activity is positively or negatively affected by any kind of external influence. Although a number of soil ecological studies considered both the active and the resident microbial community (Baldrian et al. 2012, Nunes et al. 2018, Schostag et al. 2019, Lupatini et al. 2019b), large scale mapping of shifts in the active soil microbiome is hampered by the low throughput nature and the costs of currently available kits for RNA extraction from soil. Therefore, we developed an alternative, fast and affordable method for nucleic acid extraction from soil.

The aim of this study was to investigate the resident and active microbial community in different soil- and management types and in different soil fractions (*i.e.* bulk soil and rhizosphere). To this end, we collected bulk and rhizosphere samples in two different growth stages of summer barley (*Hordeum vulgare*) in two distinct soil types – peaty and sandy - in the Netherlands, with different types of soil management. Four major organismal groups were assessed: bacteria and fungi - representing the primary decomposers - and protists and nematodes, two major grazers on the bacterial and fungal communities. Variable ribosomal DNA regions were selected for the characterization of each of the four organismal groups. We hypothesise that i) exposure to rhizosphere will have a larger effect on the active fractions of the primary decomposers (*i.e.* bacteria and fungi) than on the active fractions of representatives of the next trophic level (protists and metazoa), that ii) the impact of prolonged exposure to distinct soil management on the primary decomposer community, will reflect in comparable shifts in the active fractions of the next trophic level, protists and metazoa, and that iii) the difference in impact between the conventional soil management systems and organic management on the microbial community, will be larger than between the two conventional soil management regimes.

## Materials and Methods

### Study sites

Samples were collected from two experimental farms: (1) WUR experimental farm Vredepeel is located in the south east of the Netherlands (51°32N and 5°51E) and is characterized by sandy soil (93,3% sand, 4.5% silt, 2.2% clay) and organic matter (OM) content of 3-5%. Three different soil management strategies are applied from 2001 onwards: ConMin, ConSlu and Org. ConMin fields solely received mineral fertilizer and processed organic fertilizer without organic matter (liquid mineral concentrates), and ConSlu fields were supplemented with mineral fertilizer and slurry (pig/cow). In case of organic soil management farm yard manure, cow slurry were applied, and no pesticides were used. The six-year crop rotational system included the following major crops: potato, barley, pea, leek, maize and sugar beet, and main crops were followed by a cover crop. For further details of the set up and layout of the soil management experiments see the research report de(de Haan et al. 2018a, de Haan et al. 2018b, Quist et al. 2016, Schrama et al. 2018). (2) WUR experimental farm ‘t Kompas in Valthermond is situated in the north east of the Netherlands (52°50′N, 6°55′E) and characterized by sandy peat soil (90% sand, 7% silt, 3% clay) and a highly and variable OM content (6-14%). At Valthermond, the effect of the application of compost was investigated (yearly application of 15 tons (green) compost per hectare) in a four-year crop rotation: potato – sugar beet – potato – barley since 2013. For this study, we made a comparison between the control and compost plots.

### Soil sampling

At both sites, barley (*Hordeum vulgare*) is one of the main crop in the crop rotation system. Due to small latitudinal temperature differences, development of the barley plants in Valthermond was one week delayed as compared to the Vredepeel. Sampling was executed twice in spring 2017, during the vegetative stage and during the generative stage (Suppl. table S1).

At the Vredepeel, each of the three field was divided in 6 subfields of 540 m^2^ (Suppl. Fig. S1). In each subfield, a bulk soil and a rhizosphere sample was taken. Rhizosphere composite samples were taken by harvesting all barley plants of approximately 20 × 20 cm. Excessive soil was removed by shaking the plants and whole plants were transported to the laboratory at the field site. Bulk soil was collected by combining three individual cores in approximation of the rhizosphere sampling spot. This was done in between the barley rows with use of an auger (ø1.5 cm, depth approximately: 15 cm). In total 36 samples (18 rhizosphere and 18 bulk) were taken in each time point.

At the field laboratory, the remaining soil that adhered to the roots was brushed off from 10 individual barley plants. Rhizosphere soil and bulk soil samples were frozen in liquid nitrogen and transported on dry ice to the laboratory to be stored at −80 °C until further nucleic acid extraction.

In Valthermond we were allowed to sample in the first 2 meters of the subfield. In total 4 subfields of each treatment (suppl. Fig. S2) were sampled resulting in 16 samples (8 rhizosphere and 8 bulk) at each time point. Sampling of the barley rhizosphere was similarly executed as described earlier this paragraph. Resulting in a total of 104 samples for further analysis.

### DNA/RNA extraction and cDNA synthesis

Both DNA and RNA were simultaneously extracted from soil samples, using an in-house protocol based on phenol-chloroform-isoamylacohol extraction (see Table 1). Quality and quantity of the obtained RNA and DNA was respectively measured with a Nanodrop and Qubit. The nucleic acid eluate was stored in −80 °C upon further processing. For synthesis of cDNA from extracted RNA the Maxima First Strand cDNA Synthesis Kit for RT-qPCR (Fermentas, Thermo Fisher Scientific Inc., USA) was used according to the manufacturer’s instructions. All individual DNA and cDNA samples were diluted to respectively 1 ng/ul and 0.1 ng/ul to serve as a template for PCR amplification.

**Table 1.** Protocol for the extraction of DNA and RNA from soil.

| Step | Procedure |
| --- | --- |
| 1. | Weigh 2 g of thoroughly mixed soil, transfer it to a 15 mL bead tube), and add 1.5 g of coarse silicon carbide powder (46 grit).<br>(Keep bead tubes on ice from step 1-9.) |
| 2. | Add 2.5 mL of bead solution (181 mM disodium phosphate, 121 mM guanidinium thiocyanate), 0.25 mL of lysis buffer (150 mM NaCl, 4% (w/v) SDS, 0.5 M Tris), and 0.8 mL of a 120 mM ammonium aluminum sulfate dodecahydrate solution. |
| 3. | Add 3.5 mL of phenol:chloroform:isoamyl alcohol (25:24:1, pH: 8.0, 4°C) and mix it manually to disintegrate the biphasic layer. |
| 4. | Place the bead tubes for 10 minutes on a Digital Vortex genie 2 (SI-A256) with a SI-H512 horizontal 15 ml tube holder at maximum speed (2,850 rpm) at 4 °C. Note: place no more than 4 tubes in the tube holder as this would lower the rpm. |
| 5. | Incubate the tubes horizontally at -20°C for 10 minutes. Thereafter: repeat step 4, and continue to step 6. |
| 6. | Centrifuge the tubes for 10 minutes (2,500 x g) at 4°C to separate the soil particles from the lysate. |
| 7. | Transfer 3 mL of the upper aqueous phase to a new 15 mL tube, and add 1.5 mL of ice-cold precipitation solution (an aqueous solution of 5 M NaCl, 22 mM citric acid anhydrous salt, and 29 mM trisodium citrate dihydrate) |
| 8. | Centrifuge the tubes for 10 minutes (2,500 x g) at 4°C to separate the precipitate from the nucleic acids. |
| 9. | Transfer 4 mL of supernatant to a new 15 mL tube containing 5 mL of isopropanol at room temperature (RT). Mix gently by hand for 5s and centrifuge the tubes for 15 minutes (2,500 x g) at 4°C to precipitate the nucleic acids. |
| 10. | Discard the isopropanol, and air dry the pellet for 5 minutes at RT. |
| 11. | Add 1mL binding solution (at RT) to the pellet (binding solution: an aqueous solution of 5M guanidinium thiocyanate and 30mM Tris-HCl (pH: 6.5) with 9% (v/v) isopropanol), and vortex to redissolve the pellet. |
| 12. | Load 0.5 mL of solution to a silica spin filter (DNAK1009 Miniprep RNA column, Biocomma) to bind the nucleic acids (DNA and RNA), and spin for 30s at 10,000g. |
| 13. | Discard the flow-through, add 0.75 mL of washing solution (10mM Tris-HCl, pH: 6.5), 100mM NaCl, and absolute EtOH final v/v 50%) to the spin filter, spin for 30s at 10,000g. Repeat this step 3 times. |
| 14. | Air-dry the filter for 5 minutes, and transfer the spin filter to a clean collection tube. |
| 15. | Add 0.2 mL of elution buffer (10mM Tris-HCl, pH: 8.0) to the spin filter, and spin for 30s at 10,000g. |
| 16. | Collect the eluate (0.2 mL) and store it at -80 °C until further use. |

### PCR amplification and sequencing

The variable V4 region of bacterial 16S rRNA gene was utilized as targets for the analyses of Illumina 16S rDNA sequencing and the V9, V7-V8, V5-V7 were utilized as targets for protozoa, fungi and metazoan 18S rDNA sequencing, respectively. To prepare the samples for sequencing a twostep PCR procedure was followed. In the first PCR step, 3µl of diluted DNA or cDNA template was used to attach a locus specific primer (Table 2) to the amplicon, extended with a read area and an Illumina overhang adapter using the following temperature profile: 3 min 95°C, followed by 35x (95°C, 10 s; 55 °C, 20 s; 72 °C, 20 s) and a final extension step of 72°C of 5 min. This was done in triplicate for all samples and for each of the four organismal groups. The second PCR step was performed on 40x diluted amplicons of PCR step 1. This PCR 2 was conducted to attach the Illumina index and the Illumina sequencing adaptor (3 min 95°C, followed by 10x (95°C, 10 s; 60 °C, 30 s; 72 °C, 30 s) and a final extension step of 72°C of 5 min). Products of PCR 1 and 2 were randomly checked on gel to ensure amplification was successful. Finally, all PCR products were pooled and sent for sequencing (Bioscience, Wageningen Research, Wageningen, The Netherlands) using the Illumina MiSeq Desktop Sequencer (2*300nt paired-end sequencing) according to the standard protocols.

**Table 2:**
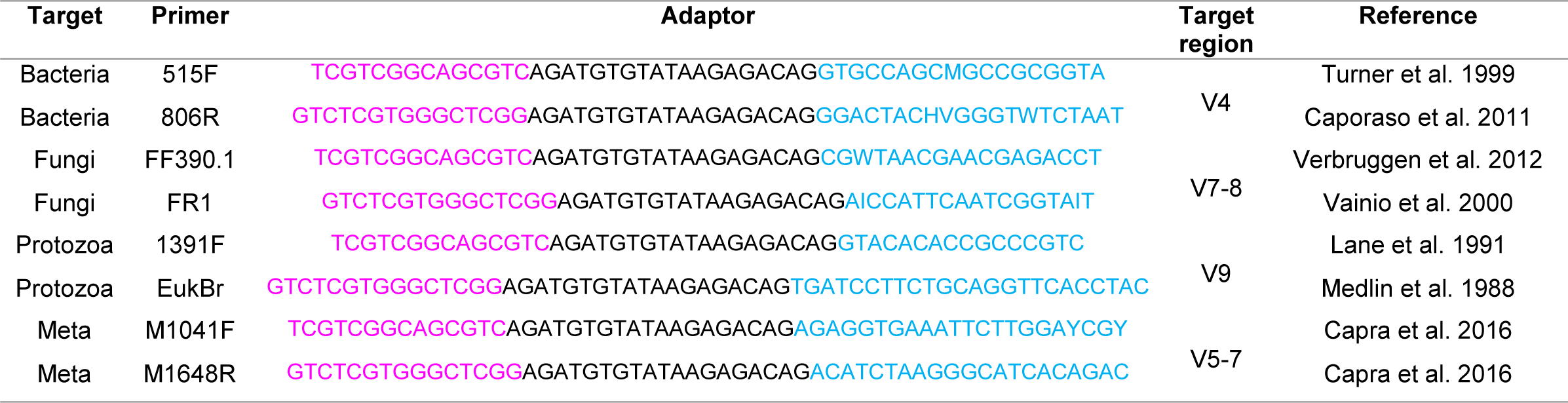
PCR1 primers - Locus-specific part (light blue), read area (black), and adaptor sequences (pink)

### Bioinformatic analysis

The composition of microbial communities of the soil samples were analysed based on the sequencing data obtained from the Illumina MiSeq platform. Reads were first sorted into the experimental samples according to their corresponding indexes combination. Thereafter, they were sorted into the four organismal groups based on their locus specific primer (General run statistics can be found in suppl. Table S2).

Sequences were processed with Hydra pipeline version 1.3.3 (Hollander 2018) implemented in Snakemake (Köster & Rahmann 2012). Forward and reverse reads were paired only for bacteria and fungi while single-end sequences were analysed for protozoa and metazoan. The four taxonomical groups were quality trimmed by BBDUK and then merged via VSEARCH (Bushnell 2015, Rognes et al. 2016). Resulting unique sequences were then clustered at 97% similarity by using the usearch_global method implemented in VSEARCH and a representative consensus sequence per de novo OTU was determined (Rognes et al. 2016). The clustering algorithm also performs chimera filtering to discard likely chimeric OTUs with UCHIME algorithm in de-novo mode (Edgar et al. 2011) implemented in VSEARCH. Sequences that passed quality filtering were then mapped to a set of representative consensus sequences to generate an OTU abundance table. Representative OTU sequences were assigned to a taxonomic classification via BLAST against the Silva database (version 12.8) for bacteria, fungi, and metazoan and PR2 database (Guillou et al. 2013) for protozoa using SINA (Pruesse et al. 2012). Sequences belonging to chloroplasts, cyanobacteria and mitochondria were discarded from the bacterial dataset; sequences not belonging to Fungi and Metazoa were removed for 18S Fungi and Metazoa datasets, respectively and Streptophyta, Metazoa, fungal and unclassified Opisthokonta sequences were filtered for Protozoa dataset. Low-abundance OTUs (those with abundance of <0.005% in the total data set) were discarded (Bokulich et al. 2013) prior to analysis. Samples were transformed using Hellingers’transformation for all downstream analyses.

### Statistical analysis

Sampling effort was estimated by Good’s coverage (Good 1953). For statistical analysis, we explored β diversity patterns by performing principal coordinate analysis (PCoA) with bray-curtis dissimilarity using QIIME software (Caporaso et al. 2012). Permutational multivariate analysis of variance (PERMANOVA) was used to compare the microbial community structure between soil managements taken from different sites and with different plant growth stages for active and resident community for 4 different taxa. This was performed with 999 permutations using the adonis function, based on Bray-Curtis distances using the “vegan” package (Oksanen et al. 2016) in R. To investigate the indicator taxa involved in the differences between resident and active community, a linear discriminate analysis (LDA) effect size (LEfSe) was conducted in Microbiome Analyst (Dhariwal et al. 2017) to explore the differential microbial populations at the family level for the four different taxa (Segata et al. 2011). A significance level of α ≤ 0.05 was used for all biomarkers evaluated in this study.

## Results

The long-term impact of various types of soil management on four major soil organismal groups was monitored in experimental field where barley was grown as main crop. The bacterial and fungal communities were mapped as main primary decomposers, whereas protists and Metazoa (mainly nematodes) were included as representatives of the next trophic level. For each the four organismal groups the resident and the active community was characterized in (1) bulk and rhizosphere soil, (2) from two soil types (sandy and peaty), (3) with five types of treatments/soil management, and (4) at two time points representing the vegetative and the generative growth stage of barley.

### General analyses of the sequencing data

A home-made protocol was used (for work flow see Table 1) to extract nucleic acids (total DNA and RNA) from 104 bulk soil and rhizosphere samples. MiSeq sequencing of organismal group-specific 16S (bacteria) or 18S (fungi, protists, and Metazoa) ribosomal DNA and cDNA fragments resulted in ≈ 31 million reads (15.5 million forward and 15.5 million reverse), and on average ≈ 75,000 reads per sample.

After filtering, a total of 8.297,203 sequences were retained comprehending 724 samples for all taxa together. Comprehensive sampling of the microbial community was obtained for all treatments, with average sequence coverage of 63%, 70%, 96% and 97% for bacteria, protozoa, fungi and metazoan, respectively determined by Good’s coverage estimator (suppl. Table S3).

### Difference in resident *versus* active communities

To gain insight in the differences between soil communities we performed a PERMANOVA for all four of the organismal communities to indicate how much of the variation is explained by a specific variable. The main findings are presented in table 3. Our analysis identified Nuclei Acid (e.g. rRNA and rDNA) as the main factor responsible for the discrepancy between samples for all four organismal groups, as it explains 14 to 25% of the overall variance (P < 0.01). The next most important explanatory factor was soil type. Location Vredepeel is characterized by sandy soils, whereas peaty soil typifies the second location (Valthermond). For the primary decomposers, a higher percentage of the overall variation (11% and 13% for bacteria and fungi) was explained soil type than for the representatives of the next trophic level (8 and 6% for protists and Metazoans) (for all groups P< 0.01).

**Table 3:** Summary of the PERMANOVA. This analysis tests differences in quantitative taxonomic composition of Bacteria, Fungi, Protozoa and Metazoa assemblage considering Nucleic Acid (cDNA/DNA), Location (Vredepeel/Valthermond), Sample type (Bulk/Rhizosphere), Treatment (ConSlu, ConMin, Org, Comp and No-Comp) and Time point (Vegetative/Generative) as factors. Differences are considered significant if P <0.01. P = probability associated with the Pseudo F statistic.

| Source | F | R2 | P |
| --- | --- | --- | --- |
| **Bacteria** |  |  |  |
| Nucleic Acid | 67.659 | 0.219 | 9.99-05 |
| Location | 32.604 | 0.106 | 9.99-05 |
| Treatment | 5.465 | 0.053 | 9.99-05 |
| Sample Type | 7.158 | 0.023 | 9.99-05 |
| Time Point | 3.700 | 0.012 | 0.00240 |
| Residuals |  | 0.410 |  |
| **Fungi** |  |  |  |
| Nucleic Acid | 48.045 | 0.144 | 9.99-05 |
| Location | 43.315 | 0.130 | 9.99-05 |
| Treatment | 11.086 | 0.100 | 9.99-05 |
| Sample Type | 7.174 | 0.022 | 9.99-05 |
| Time Point | 6.923 | 0.021 | 9.99-05 |
| Residuals |  | 0.410 |  |
| **Protozoa** |  |  |  |
| Nucleic Acid | 73.162 | 0.208 | 9.99-05 |
| Location | 27.792 | 0.079 | 9.99-05 |
| Treatment | 6.344 | 0.041 | 9.99-05 |
| Sample Type | 7.550 | 0.022 | 9.99-05 |
| Time Point | 4.788 | 0.018 | 9.99-05 |
| Residuals |  | 0.461 |  |
| **Metazoa** |  |  |  |
| Nucleic Acid | 76.172 | 0.245 | 9.99-05 |
| Treatment | 6.014 | 0.058 | 9.99-05 |
| Location | 17.300 | 0.056 | 9.99-05 |
| Sample Type | 12.655 | 0.041 | 9.99-05 |
| Time Point | 6.981 | 0.022 | 9.99-05 |
| Residuals |  | 0.429 |  |

In addition to the PERMANOVA, we generated a principal coordinate analysis (PCoA) ordination of a Bray-Curtis dissimilarity matrix (Fig. 1). The effect of Nucleic Acid (rRNA (= cDNA) for the active, and rDNA for the resident community) is clearly visible. The resident communities (blue and light blue) cluster, and are separated from the active microbial communities (red and ochre) for all four organismal groups. This separation between clusters was most obvious for Bacteria (Fig 1A) and Protists (Fig 1C). For Fungi (Fig 1B) and Metazoans (Fig 1D) this difference was significant as well, but this effect was less pronounced.

**Figure 1:**
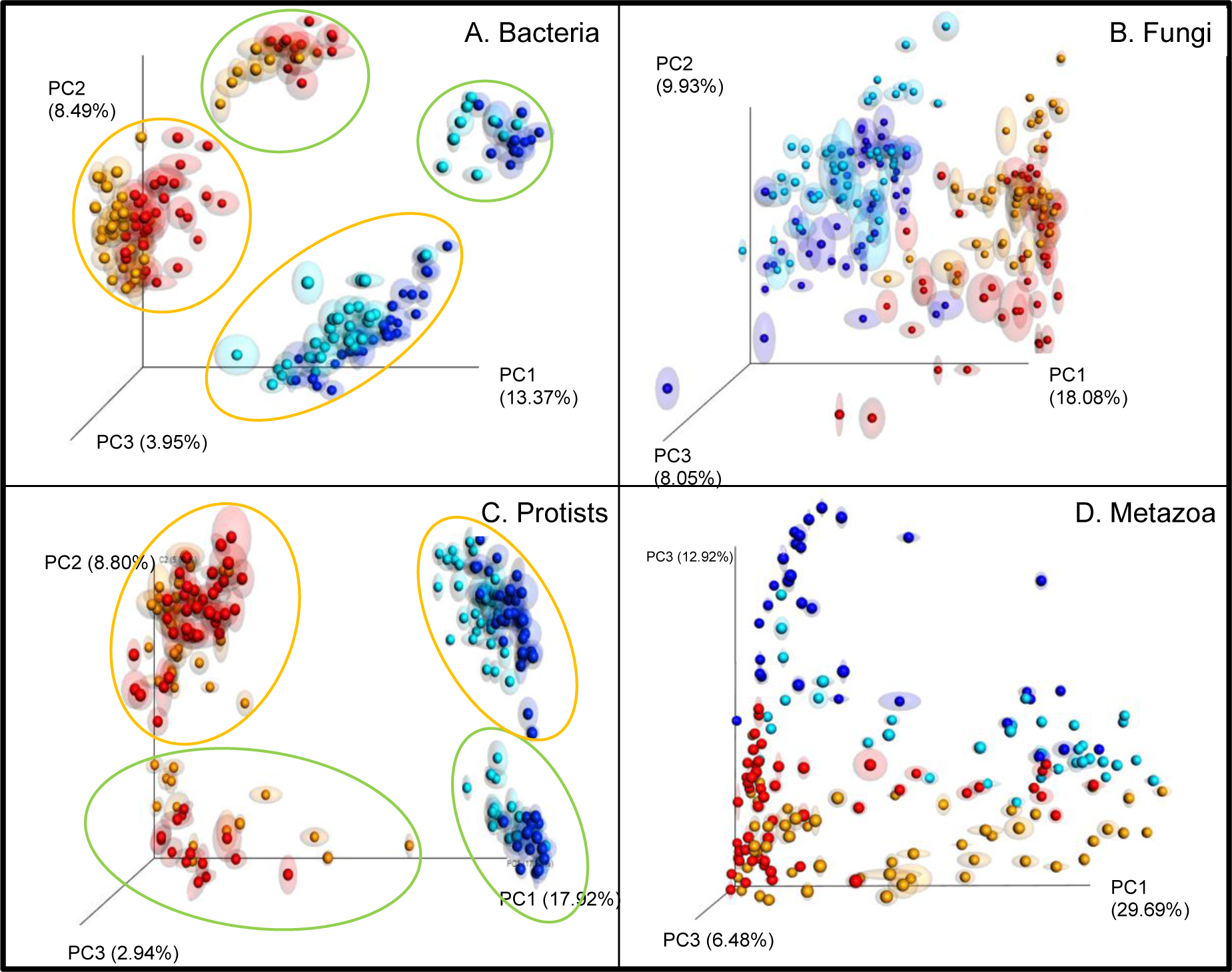
Principal coordinate analysis (PCoA) plot with Bray-Curtis dissimilarity. Plots illustrating distances between communities in all individual samples (n=208) for A: Bacteria, B: Fungi, C: Protozoa and D: Metazoa. Distinguishing between cDNA-bulk (red), cDNA-rhizo (ochre) and DNA-bulk (dark blue) DNA-rhizo (light blue). Locations are indicated by either an ochre circle (Vredepeel, sandy soil) or a green circle (Valthermond, peaty soil)

In Fig. 1, the impact of ‘Location’ is readily observable. In case of bacteria and protists, samples from Valtermond (peaty soils) are encircled in green, whereas the samples from Vredepeel (sandy soils) are defined by an ochre line. The difference between the two locations should be attributed mainly to the difference in soil type. Location effects were also observed for the fungal and metazoan community, but limitations in terms of dimensions and distinguishable colours hampered a clear visualisation of this effect (suppl. Fig S3).

Exposure to barley-induced rhizosphere conditions resulted in a shift in both the resident and the active microbial community. For bacteria, the ‘Sample Type’ effect, *i.e.* the difference in community composition between bulk and rhizosphere soil, is reflected in shifts at both DNA and cDNA level (blue and light blue, and red and ochre, respectively) (Fig. 1A). For protists, this phenomenon is most pronounced at DNA level (Fig. 1C), and at cDNA level only for the peaty soil (location Valthermond). It is noted that PERMANOVA pinpointed significant sampling type effects for all for organismal groups.

### Microbial taxa contributions to differences between resident and active communities at location Vredepeel (sandy soil)

A clear contrast in microbial community was observed between the two locations investigated in this research (Fig. 1). For pinpointing the differences between the resident and the active communities, we focussed on a single location, Vredepeel. Linear discriminant analysis (LDA) effect size (LEfSe) tests with threshold values of ≥ 2 or ≤ −2 were applied to identify specific microbial taxa that contributed to the observed differences in resident and active microbial communities. A total of 9 bacterial, 8 fungal, 10 protist and 25 metazoan orders contributed significantly to the difference between the resident (rDNA-based) and active (rRNA-based) community (Fig. 2)

**Figure 2:**
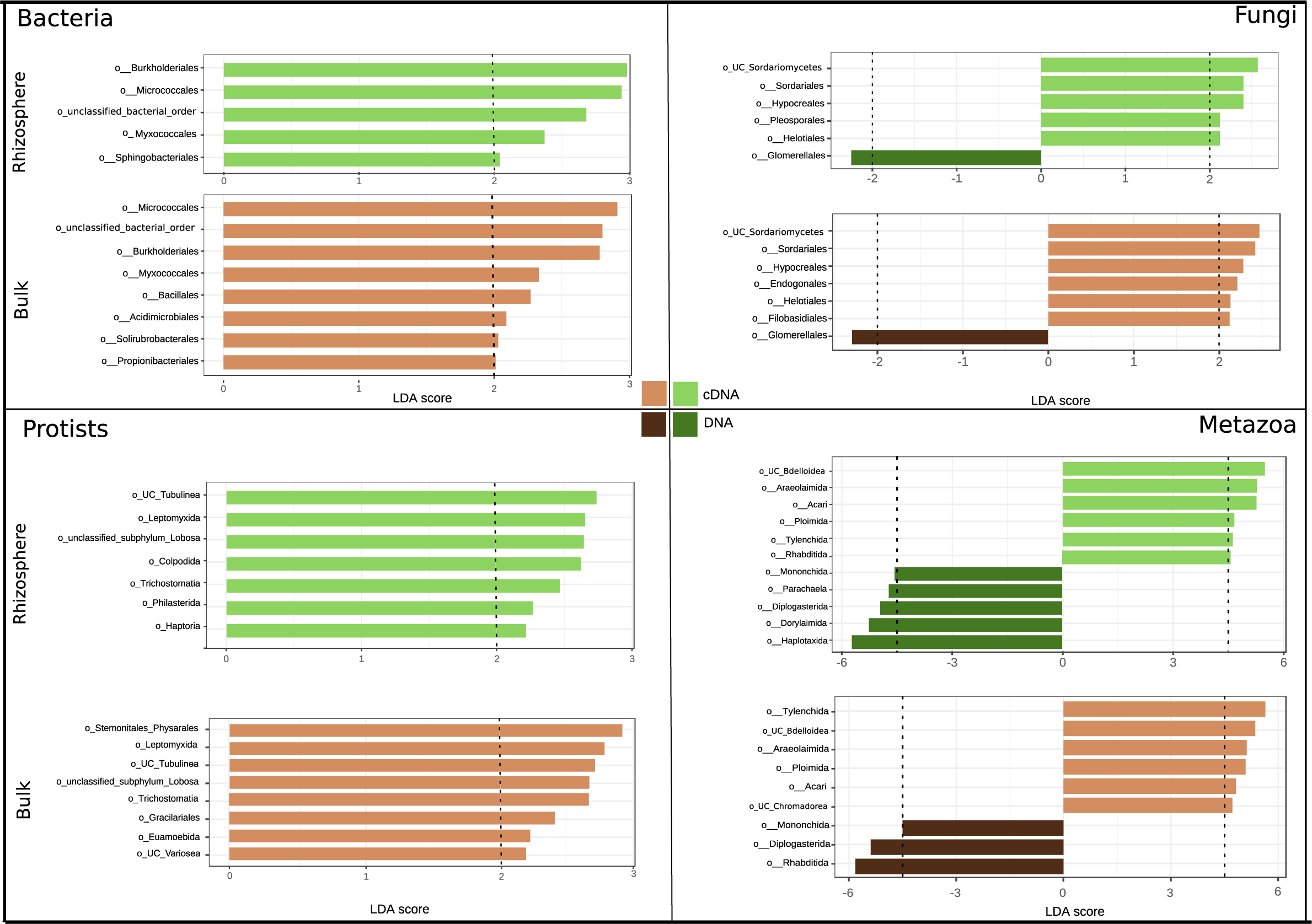
Linear discriminant analysis Effect Size (LEfSe) of bacterial, fungal, protists and metazoan OTUs, which most likely explain differences between the discrepancy in nucleic acid type (DNA or cDNA) in rhizosphere (green) or bulk soils (brown). Taxa with a ratio higher than 2 are considered active (light green and brown) and taxa with ratios lower than −2 are considered dormant (dark green and brown)

For bacteria, four orders were identified that were highly active both in rhizosphere and bulk soil (Fig. 2A). The Sphingobacteriales stood out as being highly active the barley rhizosphere, whereas Bacillales, Acidomicrobiales, Solirubrobacterales and Proprionibacteriales were predominately active in the bulk soil. It is noted only the latter order showed a large contrast with the rhizosphere community, the other three bacterial orders had LDA scores just below 2 (Data not shown).

Analysis of the fungal community revealed four orders that were highly active in both the bulk soil and the rhizosphere. The order Glomerellales, was shown to be present (DNA) but barely active in both soil compartments, hence an LDA score of −2. The bulk soil was typified by active members of the Filobasidiales and Endogonales. Notably, the contrast between bulk soil and rhizosphere was subtle for Filobasidiales, and more substantial for members of the Endogonales.

Protists, being a predominantly bacterivorous group in soil, were included as major representatives of the next trophic level. Haptoria, Philasterida and Colpodida were identified as protist orders with an enhanced metabolic activity in the barley rhizosphere. Being an extremely broad organismal group, it was no surprise to see a large number to be highly active Metazoa in the vicinity of barley roots. Our analyses revealed active rotifers, mites, nematodes, and insects in the rhizosphere compartment. Striking is the difference between the activity of the bacterivorous nematode order Rhabditida in bulk and rhizosphere soil. Rhabditida are known as extreme opportunists (Bongers 1990). This ecological characteristic is nicely reflected in Fig. 2. Limited attention will be paid to the Metazoa as they were present in relatively low numbers in the 2 g soil samples analysed in this study.

### Effects of different fertilization regimes on community structures

As compost treatment on the location Valthermond characterized by peaty soil revealed no significant soil management effects for any of analysed the organismal groups (see suppl. Table S4), we decided to focus on the impact of soil management on sandy soil location (Vredepeel). To identify the effect of the three soil management regimes, ConMin, ConSlu and Org, a PERMANOVA and a PCoA was performed on solely the sandy soil samples (see Table 4, Figure 3).

**Table 4:**
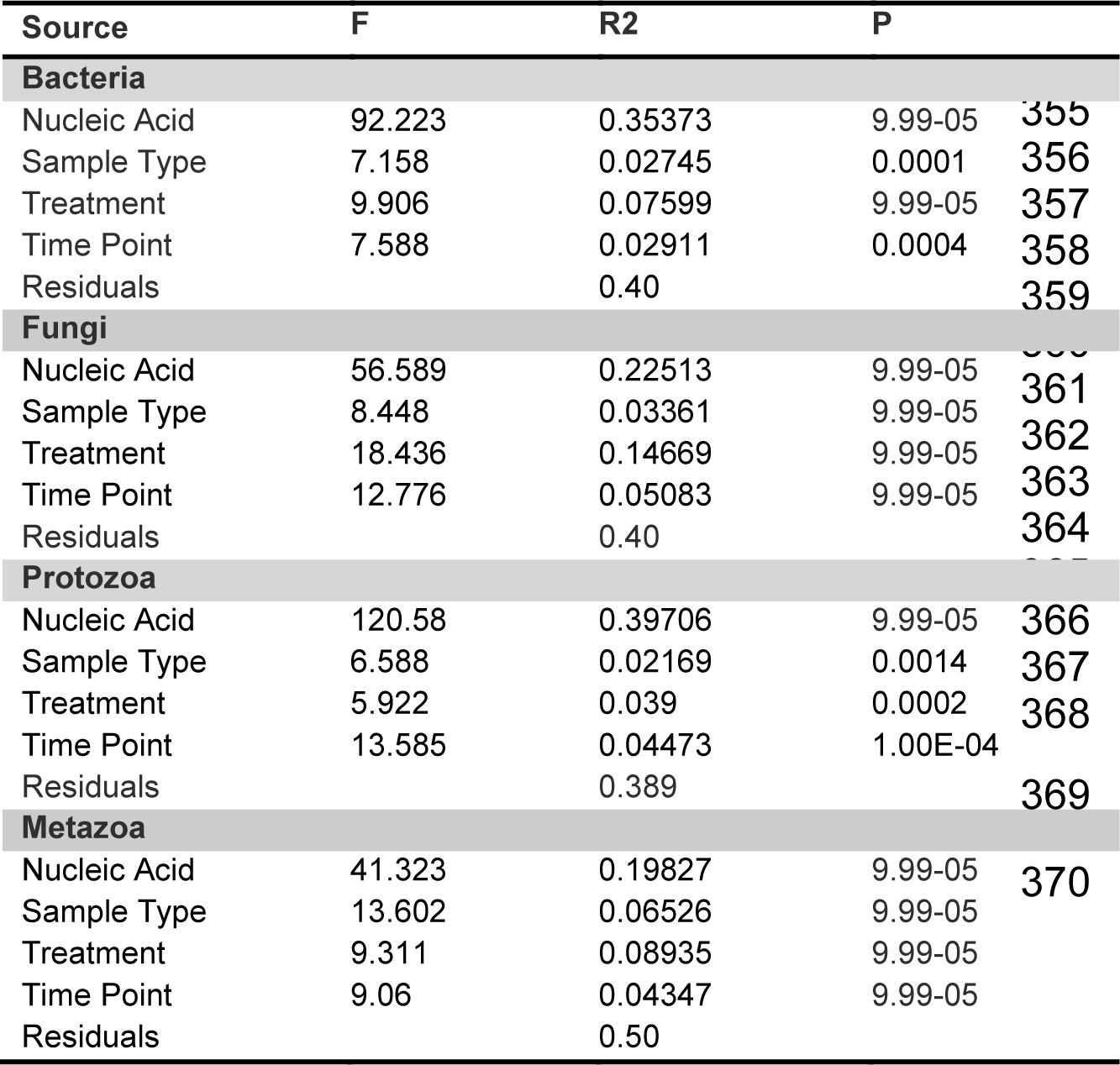
Results of PERMANOVA testing samples of Vredepeel for the effects of nucleic acid (cDNA and DNA) soil management (ConMin, ConSlu, ORG), sample type (bulk soil and rhizosphere) and time point (vegetative and generative) on soil microbial community.

**Figure 3:**
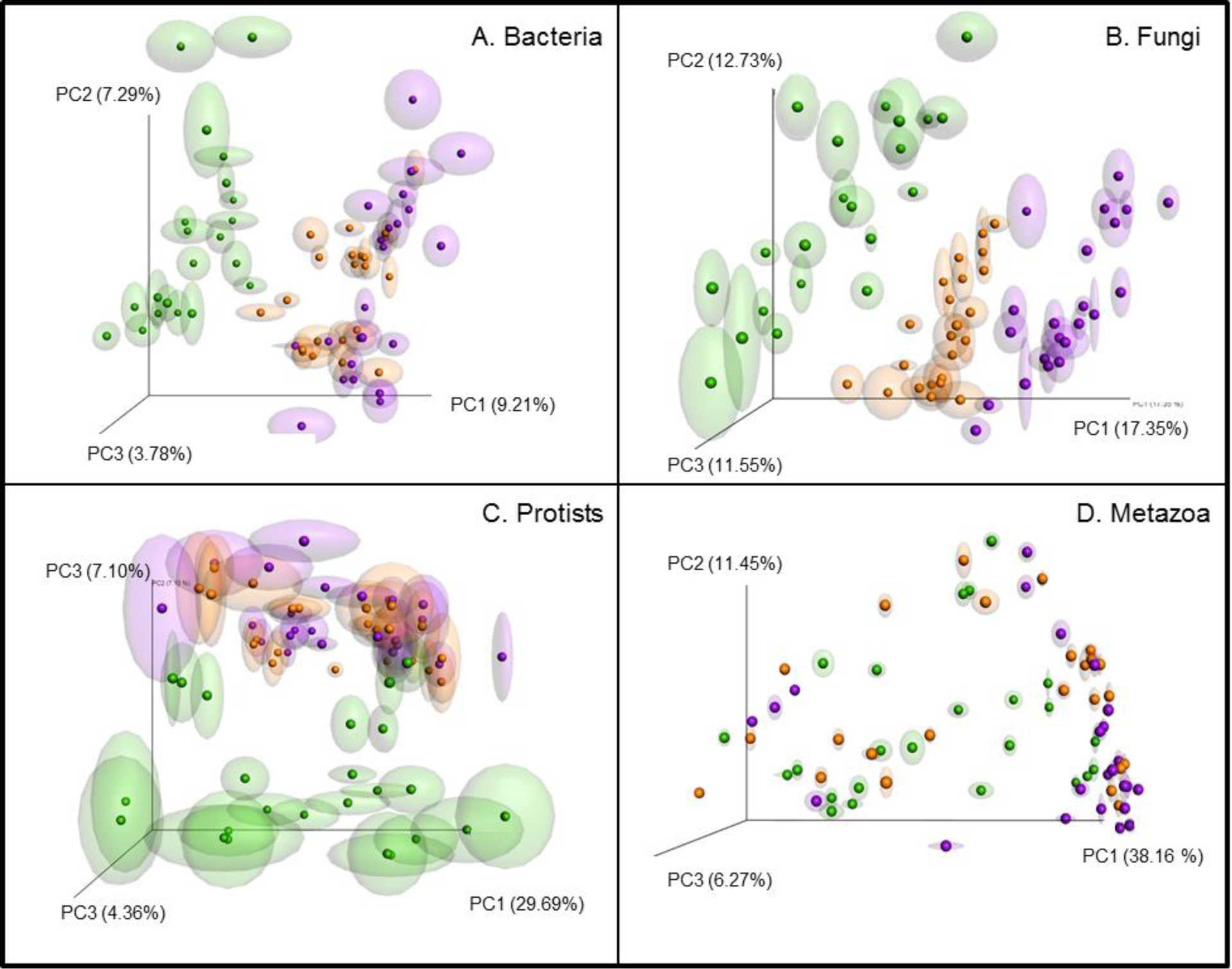
Principal coordinate analysis (PCoA) plot with Bray-Curtis dissimilarity. Plots illustrating distances between communities in all individual cDNA samples from Vredepeel (n=72) for A: Bacteria, B: Fungi, C: Protozoa and D: Metazoa. Distinguishing between treatments: ConMin (purple) ConSlu (orange) and Organic (green).

Both analyses gave a similar overall result: the microbial community structure from the Org fields was distinct from the communities found under ConMin and ConSlu soil management. This soil management effect was most obvious for Bacteria, Fungi and Protozoa. In Fig. 3, the impact of contrasting soil management regimes is shown regarding the active communities. A similar analysis on the resident community revealed a comparable pattern (suppl. Fig. S4). For Metazoans, no clear soil treatment effect was observed. In general, organic soil management increased the total OTU abundances from bacteria, fungi and protists. This was not observed for metazoans.

#### Bacteria

In general, prolonged organic soil management on sandy soil boosted the abundance of almost all microbial organisms at rDNA level (suppl. Fig. S5.1b). Out of the 38 bacterial taxa that were significantly different in terms of abundance between ConMin and Org fields, 36 taxa were significantly upregulated in the organic treatment. When considering the active fraction of the bacterial community, more taxa were found to be different. Out of the 47 significantly soil management affected bacterial taxa, 31 were more abundant in Org, while 16 showed higher activity in ConMin fields.

In the LEfSe analysis of bacterial cDNA sequences (Fig. 4) bacterial orders which active fraction was enlarged as a result of with the three soil management treatments are presented. As compared to more conventional soil treatments, prolonged organic soil management has boosted a range of bacterial orders (LDA score > 2). Desulfuromonadales, Clostridiales, Erysipelotrichales, Rhodocyclales, Rhodobacterales and Nitrosomanadales had the highest LDA scores, and their increased activity was confirmed by ANOVA (grey arrows in Suppl. Fig. S5.1). For the conventional soil treatments (ConSlu and ConMin), orders Bacillales, Deinococcales, Micrococcales, Acidobacteriales, Kineosporiales, and Streptomycetales were identified as most significant indicator taxa. ANOVA did not confirm the status of the order Kineosporiales as indicator taxon for the ConMin treatment (red arrow 4 in Suppl. Fig. 5.1a). It is noted that taxa without a formal systematic name were not taken into consideration.

**Figure 4:**
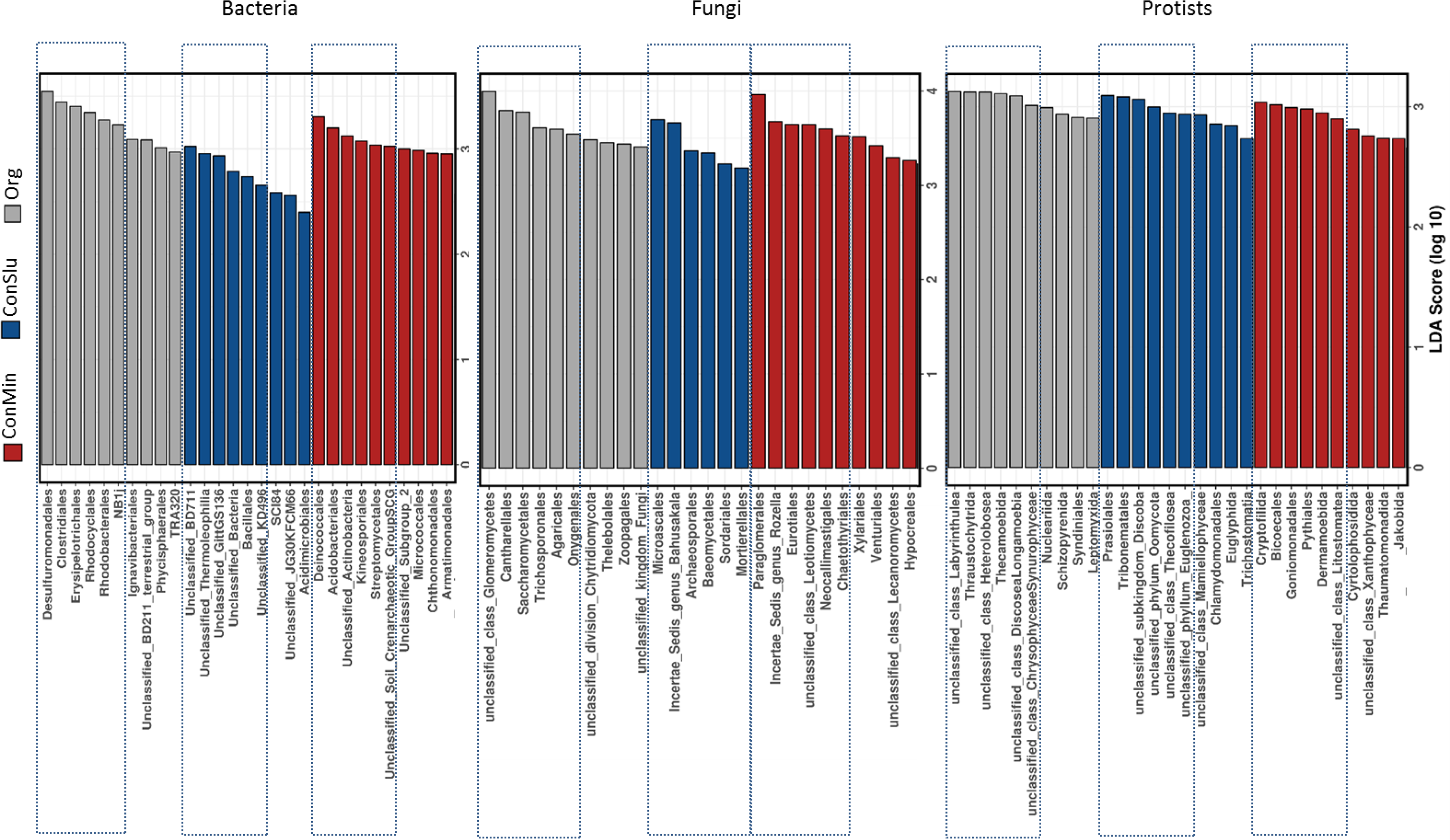
Linear discriminant analysis Effect Size (LEfSe) for differentially active bacterial, fungal, and protozoan orders between the different soil management types (ConMin=red, ConSlu=blue and Org=grey). The six most indicative orders for each treatment are delineated.

#### Fungi

Based on OTU abundances, the number of fungal orders that were promoted in organic fields was higher (22 taxa) than the number of taxa showing an increased abundance in conventional fields (8 taxa) (ConMin and ConSlu - Suppl Fig 5.2b). When concentrating on the fungal orders which activity was affected by soil management type, 19 taxa were more active in organic soils and 10 were promoted under conventional soil management regimes (ConMin and ConSlu) (Suppl Fig. 5.2a).

LEfSe analysis of fungal rRNA sequences revealed the orders Glomeromycetes (unclassified class), Cantharellales, Saccharomycetales, Trichosporonales, Agaricales, and Onygenales as most significant indicators under the organic regime (Fig. 4). In many cases, ConSlu-promoted taxa were also activated under the ConMin regime. However, the order Microascales was revealed as a specific marker for ConSlu-treated fields specifically. ANOVA confirmed the indicator status of this order for the ConSlu treatment (blue arrow 1 in Suppl. Fig. 5.2a). The orders Paraglomerales, Eurotiales, Neocallimastigales, and Chaetothyiales were identified as indicator taxa for conventional farming system using mineral fertilizers only.

#### Protozoa

The total abundance of most protists OTUs is upregulated in organic treatment as compared to conventional soil management. Comparison of significant differences between organic *versus* conventional treatment (ConMin) revealed an Org-based increase for 28 taxa, whereas 13 protozoan orders showed higher abundances under ConMin. The impact of organic soil management at RNA level showed an opposite trend. Of the 41 soil management-affected protozoan orders, only 12 taxa showed a higher OTU abundance in the fields under organic management.

Thecamoebida were identified as protist indicator taxa for organic soil management. Unclassified members of the classes Labyrinthulea and Heterolobosea were characterized by higher densities and higher activity in Org fields (Fig. 4., Suppl Fig. S 5.3a).

Prasiolales, Tribonematales, Cryptofilida, Phytiales, Dermamoebida and Bicoecales were identified as indicator taxa for conventional soil management (ConMin and ConSlu) (Fig. 4). Notably, ANOVA did not be confirmed the indicator status of Tribonematales and Bicoecales.

#### Metazoa

As compared to the bacteria, fungi and protists, the impact of the three soil managements regimes had little impact on the metazoan communities. Only a few orders were identified as marker taxa contributors by LEfSe analysis. Mononchida, an order of predatory nematodes, and the mollusc order Nuculoidea were more abundant under organic farming. The nematode orders Dorylaimida and Areaolaimida were more active under the ConSlu regime, while Tylenchida and Monhysterida were indicators for ConMin. Additionally, ANOVA gave non-corresponding results for a number of the aforementioned orders.

## Discussion

From a crop production point-of-view-intensive agricultural practices are efficient, but as an unwanted side effect these practices have contributed to a reduction of soil biodiversity and a simplification of soil food webs (Tsiafouli et al. 2014). Organic soil management is supposed to be more compatible with the preservation of soil biodiversity than conventional management practices. However, our insight in the impact of different soil management regimes is fragmented and rather superficial.

Here, we investigated the effect of various soil management systems on primary decomposers, the bacterial and fungal community, as well as major representatives of the next trophic level, protists and metazoa. The dormant part of the microbial community is considered to be very substantial, hence to assess the impact of soil management systems it is not only important to consider multiple major organismal groups, but also to map both the resident and the active fractions within these organismal groups.

We demonstrated the relevance of taking both the active and the resident soil community into consideration. For all four organismal groups, nucleic acid type (rRNA for the active fraction, rDNA for the resident community) explained a larger part of the observed variance than location (associated with soil type), the soil management regimes, or the sampling type (bulk of rhizosphere soil). These results were facilitated by a newly developed fast and affordable nucleic acids extraction protocol. Below we will firstly discuss the nature of the difference between active biota and overall biodiversity, and subsequently we will address the large fraction of soil taxa which activity is promoted more under organic management than under one of the two conventional systems.

### Limitations on the use of rRNA and rDNA data for mapping active and resident soil biota

Using rRNA and rDNA as markers for active and resident soil biota revealed large differences between these fractions. We are aware of the potential shortcomings of this type of markers. Ribosomal RNA is a highly abundant transcript, and rDNA is a multi-copy gene for which the number of copies varies enormously between organismal groups. Among bacteria, the number of rRNA operons is low and moderately diverse; typically, bacterial genes harbour 1–15 copies (Espejo & Plaza 2018). For protists, number of rDNA copies is substantially higher. Focussing on a range diatoms and dinoflagellates, individual species were shown to harbour rDNA copy number in the range of 100 to 10,000 (Godhe et al. 2008). Recently, a genome-based estimates of the number of rDNA copies among fungal taxa showed that this could range between 14 and 1,400. This variation in rDNA copy number could not be linked to trophic preferences or other easily observable ecological characteristics (Lofgren et al. 2018). From this information, it is obvious that rRNA data from microbial communities should not be used for comparisons between organismal groups, neither should they be used for comparison of abundance or activity at high taxonomic level within an organismal group.

Dormant soil inhabitants may have ribosomes with functional rRNA’s (Nanamiya et al. 2010). Such a condition allows organisms to resume activity as soon as condition are favourable again. However, it is reasonable to assume that the transition from a dormant to a physiologically active state will be accompanied by a substantial increase in rRNA level.

Hence, the use of rRNA/rDNA data for the characterisation of active and resident microbial communities in soil has a number of inherent constraints. Nevertheless, within-taxon comparison of rDNA-based sequence data under various environmental conditions is likely to reveal robust and valuable information on the impact of these conditions.

### General differences in active and resident community

Community characterisation by either of the two types of nucleic acids revealed that dormancy is a phenomenon relevant for all four organismal groups under investigation here.

#### Bacteria

A number of highly active bacterial orders including Burkholderiales, Micrococcales and Myxococcales, were shown to be active both in bulk and rhizosphere soil. Sphingobacteriales are an exception as members of this order were far more active in the rhizosphere (Fig. 2). (Peiffer et al. 2013) found a similar enrichment of this order in the rhizosphere of maize. Next to maize, (Haichar et al. 2008) showed Sphingobacteriales accumulation on roots of wheat, rale and barrel clover. This apparently general rhizosphere accruement is in line with the observed upregulated activity near the roots of barley. Propionibacteriales were identified as being specifically active in bulk soil. Propionibacteriales are known to contribute to both primary and secondary fermentations (Johnson & Cummins 1972), and this activity could be related to the distinct fertilisation regimes. However it appears not selected in the rhizosphere of barley plants.

#### Fungi

Both in bulk soil and the rhizosphere representatives of the orders of Sordariales, Hypocreales, Helotiales were highly active (all belonging to the Ascomycota). This in accordance with a survey on four conventional arable fields in Austria agricultural that revealed the dominant presence of the same three fungal order (Klaubauf et al. 2010). However, the high abundance of these Ascomycete orders is not specific for arable soils, (Tedersoo et al. 2009) found similar orders associated with tree roots. Both studies are based on DNA data, and our data demonstrate that these fungal orders are highly active. Glomerellales (Ascomycota) showed a negative LDA score for bulk as well as rhizosphere, indicating dormancy. Drought has been shown to specifically decrease protein abundances of Glomerellales (Bastida et al. 2017). As sampling took place during a severe period of drought, this could be a clarification of the observed shift towards the DNA site of the spectrum. Unfortunately, very little is known about the ecology of Glomerellales.

#### Protozoa

Barley rhizosphere was characterized by the presence of active representatives of the orders of Haptoria, Colpodida and Philasterida. Haptoria are ubiquitous free-living predatory ciliates in soils (Vd’acny et al. 2014), whereas Copodida are predominantly grazers of bacteria (Vdacny & Foissner 2018). The high bacterial density in the rhizosphere could be an explanation for the accumulation of active Copodida. Philasterida are known to occur in terrestrial habitats, but barely any information is available about their ecology. Bulk was typified by active Euamoebida, and this was also found by (Geisen et al. 2015) in grasslands and forest mineral soils. In the same study, the paraphyletic class Variosea was found as a dominant active class in bulk soil collected from a grassland. We could confirm the presence of active members of this class in bulk soil from barley fields. This class was defined in 2004 (Cavalier-Smith et al. 2004), and little is known about their ecological role in soil. Physarales is suggested to play an important role in litter breakdown (Kamono et al. 2009) and therefore it is no surprise that they are specifically active in bulk soils.

#### Metazoa

As compared to Bacteria, Fungi and Protists, Metazoa showed the strongest compartment effect (R^2^ = 0.041, Table 3 (Metazoa / Sample type). This indicates a high difference in community composition between bulk soil and rhizosphere. The highly density and activity of soil micro fauna has been well documented in a range of systems (Chen et al. 2007, Griffiths 1994, Irshad et al. 2011). Our analysis revealed also a number of orders for which a relatively high fraction was dormant. This is unexpected as for instance member of the speciose nematode order Dorylaimida are expected to be non-dormant in the rhizosphere. In a more generalized manner, Hu et al (2016) pointed at the same phenomenon: metazoa are abundant in rDNA samples while a relatively low number of rRNA sequences is detected. A potential explanation for this observation could be relatively low number of RNA copies relative to DNA in metazoans (Hu et al. 2016).

### Impact of soil management practices on the rhizobiome

The effect of organic soil management on bacterial communities is relatively well documented (Hartmann et al. 2015, Lori et al. 2017, Lupatini et al. 2017, Suleiman et al. 2018). To the best of our knowledge there are no other studies investigating the effects of distinct soil management regimes on four organismal groups simultaneously.

#### Bacteria

At DNA level, ANOVA revealed that 44 out of the 48 bacterial taxa were more abundant under organic management (Suppl. Fig. 5.1b). Regarding bacteria, this observation is in line with earlier research the effects of organic soil management (Mäder et al. 2002) (e.g.). However, at activity level (as revealed by rRNA-based analysis) another picture arose: ‘only’ about 1/3 of the 50 bacterial taxa were most active under organic soil management conditions (Suppl. Fig. 5.1a). As microbial activity matters in terms of soil food web functioning, these results underline the relevance of taking – next to abundance data – activity data into account.

The orders that seem to be the most contributing to the difference in organic *versus* conventional soil management (in terms of abundance OTU and LEfSe score) in the active community were Desulfuromonadales (δ-proteobacteria), Clostridiales (Firmicutes), Rhodocyclales (β-proteobacteria), Rhodobacterales (α-proteobacteria). Desulfuromonadales were a typifying order in Org fields. This order harbours a range of sulphate and sulphur reducing bacteria (Warren et al. 2016). There enhanced activity could relate to the slightly higher S content of the Org fields (on average 247 mg S/kg soil, ConMin and ConSlu: 193 and 214 mg S/kg over period 2011 – 2016)). Clostridiales are metabolically diverse, however it has been shown that it increase with the addition of organic matter with a high recalcitrant C content (Goldfarb et al. 2011). Org treatment, crop residues were use as green manure (Quist et al. 2016). Rhodocyclales and Rhodobacterales play a role in denitrification (Saito et al. 2008, Yoshida et al. 2009).

Conventional fields were characterized by highly active Actinobacteria (Kineosporiales, Streptomycetales and Micrococcales). This difference was also observed in previous research on the impact of conventional and organic cropping systems. In a survey over 3 years, Orr et al (2015) detected a similar increase in Actinobacteria, but they also showed a strong sample year effect. For the interpretation of our data this should be taken into consideration.

#### Fungi

In total 70% of the significantly soil management-affected fungal orders were more abundant in Org. In general, higher abundance was accompanied by higher activity. As expected, the majority of the fungal OTUs were assigned to the Ascomycota. They are important decomposer of organic substrates (such as leaf litter, wood, and manure) and more studies reported them as the major fungal phylum present in agro-ecosystems (Francioli et al. 2016, Lienhard et al. 2014). The two Basidiomycetes (Agaricales, Cantharellales) were found to be abundant in organic fields. As they are both known to be decomposers this was expected (Floudas et al. 2015). Onygenales were rarely found in the conventional fields but highly active in the organic soils. This order is associated with animal dung (Doveri et al. 2012, Sugiyama et al. 2002). So, active members of the Onygenales are probably the result of the application of cattle manure in the Org plots.

The strongest fungal indicator for organic farming was an “unclassified class of Glomeromycetes’. Glomeromycetes are known to form arbuscular mycorrhizas and colonize the roots of vascular land plants including barley (Williams et al. 2017). In a recent study on the same experimental farm (Vredepeel), AM fungi were also found to be more abundant under organic soil management (Martinez-Garcia et al. 2018, Lupatini et al. 2019a). AM fungi can stimulate the decomposition of recalcitrant organic matter, and makes nitrogen bioavailable (Hodge 2001). Hence, the distinct type of manure used under organic management might explain the specific stimulation of Glomeromycetes.

Paraglomerales, another member of the Glomeromycota was predominantly found in conventional systems. This finding was corroborated by (Dai et al. 2014) who found *Paraglomus* to be positively associated with the conventional production of wheat and while enhanced presence of Claroideoglomus was observed under organic management. The relation between *Paraglomerales* and fertilization system would deserve further investigation.

#### Protozoa

In contrast to the primary decomposers, organic soil management decreased the activity of many protozoa. These results contrasted with the observed stimulating effect of the organic regime on total abundances (rDNA). Increased densities of protozoa as a result of organic amendments have been reported before (Treonis et al. 2010). Under controlled greenhouse conditions, application of organic fertilizers increased bacterivorous and omnivorous protists, and strongly reduced the relative abundance of plant pathogenic protists (Xiong et al. 2018). We aren’t aware of other studies on the impact of organic amendment to soil protist activity.

#### Metazoa

Organic soil management stimulated the activity an order of predatory nematodes Mononchida, and members of the mollusc order Nuculoidea. Predatory Monochida feed on other nematodes, but this does not hold for all life stages. Larval stages are too small to capture other nematodes, and they feed on bacteria (Yeates, 1987). The strongly enriched bacterial community under the organic regimes may have promoted the activity of the Monochida. The impact of soil management on Metazoa will not be discussed further, as the numbers of individuals present in the 2 g soil samples are too low (with some nematode taxa as an exception). Hence, sampling effects could easily obscure soil management effects.

## Conclusion

The mapping of the effects of soil management on bacterial, fungal, protist and metazoan communities in barley fields underlined the relevance of the parallel monitoring of both the active (rRNA-based) and the resident (rDNA) fractions. Differences in duration of exposure to a given treatment resulted in differential responses among the four organismal groups. Short exposure (months), in this case the rhizosphere effect caused by barley, was mainly reflected in the active fractions bacteria and only to a lesser extend for fungi. Regarding metazoa, no statements can be made due to the low numbers of individuals per sample. Hence, our initial hypothesis is partly correct. Secondly, we hypothesized that prolonged exposure to distinct soil management regimes (10 years) resulted in similar large shifts in the active fractions of all four organismal groups. This appeared to be true for three organismal groups only. Also here, metazoa couldn’t be taken into full consideration as their numbers per sample were relatively low. This second assumption is therefore partially correct. Thirdly, we wanted to see whether the difference between a highly conventional (ConMin) and a more integrated soil management system (ConSlu) would be smaller than the contrast between the two conventional systems and the organic regime. This shown to be correct, and was most extensively demonstrated in the rRNA and rDNA-based ANOVA analyses.

The home-made protocol for a time-efficient simultaneous extraction of RNA and DNA from soil was highly contributory in our research. For reasons pinpointed earlier in this paper, large-scale monitoring surveys of the active fractions of the microbial soil community are relatively scarce. Hence, a substantial part of the discussion of our results was necessarily based on papers about the effects of soil management on resident communities. In follow up papers, we will investigate sample year effects, crop effect and the impact of crop growth stage on the active and resident fractions of the astonishing diverse soil living community.

## Acknowledgment

We would like to thank PPO-Vredepeel and PPO-Valthermond for operation long term filed experiments on the effect of various farming systems on a broad range of agronomic parameters and the collection of all primary farm data. This research was supported by NWO Groen grants (numbers: 870.15.021 and 870.15.022).

## Author contributions

P.H., A.K.A.S., J.J.d.H, E.E.K and J.H. were responsible for the experimental design. P.H., A.K.A.S., M.H.M.H, E.E.K and J.H. were involved in sampling the rhizosphere. S.J.J.E. developed the RNA/DNA isolation protocol. P.H. tested the RNA/DNA protocol and performed RNA/DNA isolation of the soil samples. S.J.J.E. developed the PCR two step system prior to sequencing. P.H. tested and performed the two step PCR reactions in order to prepare the sequence library. M.H.M.H. sorted the sequence data. A.K.A.S. analysed the sequence data and performed the bioinformatics and statistical analysis. P.H. and J.H. wrote the manuscript; all others co-commented on the manuscript.

