## Supplementary tables 1-4 for "Mapping of long-term impact of conventional and organic soil management on resident and active fractions of rhizosphere communities of barley"

**S1**: Barley sampling dates of the two locations

| | **Vredepeel – ConMin, ConSlu, Org** | **Valthermond – Compost, Control** |
|---|---|---|
| **Vegetative stage** | 7$^{th}$ of June 2017 | 14$^{th}$ of June 2017 |
| **Generative stage** | 10$^{th}$ of July 2017 | 13$^{th}$ of July 2017 |

**S2**: General run statistics of MiSeq.

| | **Forward reads** | **Reverse reads** |
|---|---|---|
| **Average reads/sample** | 74694 | 74694 |
| **Total reads** | 15536444 | 15536444 |
| **Average bases/sample** | 17488215 | 17754199 |
| **Total bases** | 3637548813 | 3692873434 |
| **% bases Phred score > 30 (p<0.001)** | 89.6% | 76.1% |

**S3**: Number of sequences and coverage

| Taxa-Compartment | Vredepeel | | | Taxa-Compartment | Vredepeel | | |
|---|---|---|---|---|---|---|---|
| | Sample | Number sequences - singletons | Good's coverage (%) | | Sample | Number sequences - singletons | Good's coverage (%) |
| Bacteria-Bulk | 100-DNA-VR-B-ORG-4 | 30746 | 58.3 | Bacteria-Rhizosphere | 101-DNA-VR-R-ORG-5 | 28357 | 58.7 |
| | 102-DNA-VR-B-ORG-5 | 29230 | 58.8 | | 103-DNA-VR-R-ORG-6 | 27111 | 59.9 |
| | 104-DNA-VR-B-ORG-6 | 13012 | 60.8 | | 123-cDNA-VR-R-CONMIN-2 | 15810 | 59.1 |
| | 124-cDNA-VR-B-CONMIN-2 | 12887 | 60.7 | | 125-cDNA-VR-R-CONMIN-3 | 3020 | 63.1 |
| | 126-cDNA-VR-B-CONMIN-3 | 1907 | 61.8 | | 127-cDNA-VR-R-CONMIN-4 | 9567 | 62.1 |
| | 128-cDNA-VR-B-CONMIN-4 | 13366 | 61.6 | | 129-cDNA-VR-R-CONMIN-5 | 11595 | 64.6 |
| | 130-cDNA-VR-B-CONMIN-5 | 8500 | 62.5 | | 131-cDNA-VR-R-CONMIN-6 | 17784 | 66.2 |
| | 132-cDNA-VR-B-CONMIN-6 | 16317 | 60.7 | | 133-cDNA-VR-R-CONSLU-1 | 11452 | 64.1 |
| | 134-cDNA-VR-B-CONSLU-1 | 4549 | 63 | | 135-cDNA-VR-R-CONSLU-2 | 15416 | 64.1 |
| | 136-cDNA-VR-B-CONSLU-2 | 16335 | 60.2 | | 139-cDNA-VR-R-CONSLU-4 | 12619 | 61.1 |
| | 138-cDNA-VR-B-CONSLU-3 | 13386 | 64.5 | | 141-cDNA-VR-R-CONSLU-5 | 14360 | 62.1 |
| | 140-cDNA-VR-B-CONSLU-4 | 12883 | 62.6 | | 143-cDNA-VR-R-CONSLU-6 | 12789 | 60.5 |
| | 142-cDNA-VR-B-CONSLU-5 | 8974 | 64.2 | | 147-cDNA-VR-R-ORG-2 | 16687 | 61.4 |
| | 144-cDNA-VR-B-CONSLU-6 | 8904 | 64.1 | | 149-cDNA-VR-R-ORG-3 | 16007 | 58.8 |
| | 146-cDNA-VR-B-ORG-1 | 13979 | 58.8 | | 151-cDNA-VR-R-ORG-4 | 27295 | 59.1 |
| | 148-cDNA-VR-B-ORG-2 | 18130 | 58.5 | | 155-cDNA-VR-R-ORG-6 | 9630 | 62.9 |
| | 152-cDNA-VR-B-ORG-4 | 22000 | 58.9 | | 173-cDNA-VR-R-CONMIN-1 | 14481 | 63 |
| | 154-cDNA-VR-B-ORG-5 | 5215 | 60.7 | | 175-cDNA-VR-R-CONMIN-2 | 6928 | 63.6 |
| | 156-cDNA-VR-B-ORG-6 | 7793 | 60.6 | | 177-cDNA-VR-R-CONMIN-3 | 5960 | 67.3 |

| | | | | | |
|---|---|---|---|---|---|
| 174-cDNA-VR-B-CONMIN-1 | 8289 | 60.1 | 179-cDNA-VR-R-CONMIN-4 | 16259 | 61.1 |
| 176-cDNA-VR-B-CONMIN-2 | 13260 | 59.6 | 17-DNA-VR-R-CONMIN-1 | 1460 | 62.3 |
| 178-cDNA-VR-B-CONMIN-3 | 25456 | 57.7 | 181-cDNA-VR-R-CONMIN-5 | 36233 | 63.6 |
| 180-cDNA-VR-B-CONMIN-4 | 11385 | 57.9 | 183-cDNA-VR-R-CONMIN-6 | 10992 | 61.4 |
| 182-cDNA-VR-B-CONMIN-5 | 26428 | 63.1 | 185-cDNA-VR-R-CONSLU-1 | 17036 | 67.1 |
| 184-cDNA-VR-B-CONMIN-6 | 21589 | 59.7 | 187-cDNA-VR-R-CONSLU-2 | 8654 | 65.6 |
| 186-cDNA-VR-B-CONSLU-1 | 7733 | 61.9 | 189-cDNA-VR-R-CONSLU-3 | 23521 | 63.9 |
| 188-cDNA-VR-B-CONSLU-2 | 10043 | 62.9 | 191-cDNA-VR-R-CONSLU-4 | 11521 | 64.1 |
| 18-DNA-VR-B-CONMIN-1 | 2730 | 61.9 | 193-cDNA-VR-R-CONSLU-5 | 18111 | 63 |
| 190-cDNA-VR-B-CONSLU-3 | 12394 | 63.2 | 195-cDNA-VR-R-CONSLU-6 | 21904 | 61 |
| 192-cDNA-VR-B-CONSLU-4 | 14812 | 58.6 | 197-cDNA-VR-R-ORG-1 | 24534 | 59.5 |
| 194-cDNA-VR-B-CONSLU-5 | 16501 | 61.3 | 199-cDNA-VR-R-ORG-2 | 13806 | 60.3 |
| 196-cDNA-VR-B-CONSLU-6 | 14568 | 62.5 | 19-DNA-VR-R-CONMIN-2 | 1814 | 60.1 |
| 198-cDNA-VR-B-ORG-1 | 14985 | 58.1 | 201-cDNA-VR-R-ORG-3 | 3727 | 59.2 |
| 200-cDNA-VR-B-ORG-2 | 10968 | 57.4 | 203-cDNA-VR-R-ORG-4 | 28903 | 57.8 |
| 202-cDNA-VR-B-ORG-3 | 23100 | 57.1 | 205-cDNA-VR-R-ORG-5 | 26393 | 58.3 |
| 204-cDNA-VR-B-ORG-4 | 36170 | 63.2 | 207-cDNA-VR-R-ORG-6 | 18960 | 58.3 |
| 206-cDNA-VR-B-ORG-5 | 37305 | 59.9 | 21-DNA-VR-R-CONMIN-3 | 4203 | 63.3 |
| 208-cDNA-VR-B-ORG-6 | 24941 | 57.3 | 23-DNA-VR-R-CONMIN-4 | 1040 | 64 |
| 20-DNA-VR-B-CONMIN-2 | 1505 | 66.5 | 27-DNA-VR-R-CONMIN-6 | 1786 | 67.9 |
| 26-DNA-VR-B-CONMIN-5 | 2168 | 61 | 31-DNA-VR-R-CONSLU-2 | 2220 | 61.7 |
| 28-DNA-VR-B-CONMIN-6 | 1373 | 65.9 | 33-DNA-VR-R-CONSLU-3 | 1058 | 63.7 |
| 30-DNA-VR-B-CONSLU-1 | 1205 | 62 | 35-DNA-VR-R-CONSLU-4 | 1947 | 62.4 |
| 34-DNA-VR-B-CONSLU-3 | 1041 | 61.9 | 37-DNA-VR-R-CONSLU-5 | 2976 | 63 |

| | | | | | | | |
|---|---|---|---|---|---|---|---|
| | 36-DNA-VR-B-CONSLU-4 | 1107 | 62.9 | | 39-DNA-VR-R-CONSLU-6 | 1928 | 61.8 |
| | 38-DNA-VR-B-CONSLU-5 | 1234 | 65.5 | | 41-DNA-VR-R-ORG-1 | 1018 | 59.2 |
| | 42-DNA-VR-B-ORG-1 | 2552 | 61.4 | | 43-DNA-VR-R-ORG-2 | 2206 | 60.5 |
| | 44-DNA-VR-B-ORG-2 | 8051 | 60 | | 45-DNA-VR-R-ORG-3 | 1624 | 58.7 |
| | 46-DNA-VR-B-ORG-3 | 1160 | 60.1 | | 49-DNA-VR-R-ORG-5 | 1552 | 60.6 |
| | 48-DNA-VR-B-ORG-4 | 1016 | 61.8 | | 51-DNA-VR-R-ORG-6 | 1172 | 63.5 |
| | 50-DNA-VR-B-ORG-5 | 1077 | 64 | | 69-DNA-VR-R-CONMIN-1 | 1706 | 62.7 |
| | 52-DNA-VR-B-ORG-6 | 1838 | 62.3 | | 71-DNA-VR-R-CONMIN-2 | 1723 | 63.7 |
| | 70-DNA-VR-B-CONMIN-1 | 1372 | 62.5 | | 73-DNA-VR-R-CONMIN-3 | 1425 | 63.5 |
| | 74-DNA-VR-B-CONMIN-3 | 1013 | 60.8 | | 75-DNA-VR-R-CONMIN-4 | 5874 | 64.5 |
| | 76-DNA-VR-B-CONMIN-4 | 2083 | 59.8 | | 77-DNA-VR-R-CONMIN-5 | 1203 | 65.8 |
| | 78-DNA-VR-B-CONMIN-5 | 1402 | 64.2 | | 79-DNA-VR-R-CONMIN-6 | 1588 | 62.2 |
| | 80-DNA-VR-B-CONMIN-6 | 1435 | 61.4 | | 81-DNA-VR-R-CONSLU-1 | 1429 | 64.2 |
| | 82-DNA-VR-B-CONSLU-1 | 1514 | 59.2 | | 83-DNA-VR-R-CONSLU-2 | 3415 | 64.8 |
| | 84-DNA-VR-B-CONSLU-2 | 2360 | 61.3 | | 85-DNA-VR-R-CONSLU-3 | 1266 | 65.6 |
| | 88-DNA-VR-B-CONSLU-4 | 1730 | 62.2 | | 87-DNA-VR-R-CONSLU-4 | 1450 | 65.5 |
| | 90-DNA-VR-B-CONSLU-5 | 5499 | 60.6 | | 89-DNA-VR-R-CONSLU-5 | 3350 | 62.1 |
| | 92-DNA-VR-B-CONSLU-6 | 1800 | 61.1 | | 91-DNA-VR-R-CONSLU-6 | 8127 | 60.6 |
| | 94-DNA-VR-B-ORG-1 | 1410 | 56.2 | | 95-DNA-VR-R-ORG-2 | 2639 | 62 |
| | 96-DNA-VR-B-ORG-2 | 1624 | 62.8 | | 97-DNA-VR-R-ORG-3 | 12236 | 59.9 |
| | 98-DNA-VR-B-ORG-3 | 6118 | 63 | | 99-DNA-VR-R-ORG-4 | 20942 | 57.8 |
| | Total | 645457 | | | Total | 653799 | |
| Protozoa-Bulk | 100-DNA-VR-B-ORG-4 | 14765 | 70.9 | Protozoa-Rhizosphere | 101-DNA-VR-R-ORG-5 | 17803 | 61.6 |
| | 102-DNA-VR-B-ORG-5 | 13796 | 67.4 | | 103-DNA-VR-R-ORG-6 | 16852 | 64.8 |
| | 104-DNA-VR-B-ORG-6 | 16158 | 66.3 | | 121-cDNA-VR-R-CONMIN-1 | 1750 | 66.1 |
| | 122-cDNA-VR-B-CONMIN-1 | 3401 | 61.8 | | 123-cDNA-VR-R-CONMIN-2 | 49546 | 67.3 |
| | 124-cDNA-VR-B-CONMIN-2 | 77439 | 62.2 | | 125-cDNA-VR-R-CONMIN-3 | 9131 | 65 |
| | 126-cDNA-VR-B-CONMIN-3 | 7140 | 62.4 | | 127-cDNA-VR-R-CONMIN-4 | 17852 | 65 |

| | | | | | |
|---|---|---|---|---|---|
| 128-cDNA-VR-B-CONMIN-4 | 40205 | 59.7 | 129-cDNA-VR-R-CONMIN-5 | 30150 | 66.5 |
| 130-cDNA-VR-B-CONMIN-5 | 13691 | 61.8 | 131-cDNA-VR-R-CONMIN-6 | 24703 | 67.8 |
| 132-cDNA-VR-B-CONMIN-6 | 27811 | 65.7 | 133-cDNA-VR-R-CONSLU-1 | 11281 | 68 |
| 134-cDNA-VR-B-CONSLU-1 | 2863 | 60.7 | 135-cDNA-VR-R-CONSLU-2 | 8840 | 67.6 |
| 136-cDNA-VR-B-CONSLU-2 | 15315 | 62.8 | 139-cDNA-VR-R-CONSLU-4 | 27784 | 68.2 |
| 138-cDNA-VR-B-CONSLU-3 | 25322 | 61.2 | 141-cDNA-VR-R-CONSLU-5 | 16952 | 63 |
| 140-cDNA-VR-B-CONSLU-4 | 28861 | 62.5 | 143-cDNA-VR-R-CONSLU-6 | 19304 | 62.7 |
| 142-cDNA-VR-B-CONSLU-5 | 17646 | 64.5 | 145-cDNA-VR-R-ORG-1 | 2614 | 72.1 |
| 144-cDNA-VR-B-CONSLU-6 | 30281 | 63.3 | 147-cDNA-VR-R-ORG-2 | 51788 | 78.9 |
| 146-cDNA-VR-B-ORG-1 | 58112 | 70.1 | 149-cDNA-VR-R-ORG-3 | 35025 | 67.6 |
| 148-cDNA-VR-B-ORG-2 | 52759 | 62.8 | 151-cDNA-VR-R-ORG-4 | 42866 | 69.8 |
| 150-cDNA-VR-B-ORG-3 | 3110 | 65.8 | 153-cDNA-VR-R-ORG-5 | 6024 | 67.9 |
| 152-cDNA-VR-B-ORG-4 | 51433 | 61.3 | 155-cDNA-VR-R-ORG-6 | 21509 | 70 |
| 154-cDNA-VR-B-ORG-5 | 48465 | 64.4 | 173-cDNA-VR-R-CONMIN-1 | 56957 | 73.3 |
| 156-cDNA-VR-B-ORG-6 | 42672 | 61.1 | 175-cDNA-VR-R-CONMIN-2 | 65931 | 65.3 |
| 174-cDNA-VR-B-CONMIN-1 | 39938 | 63.1 | 177-cDNA-VR-R-CONMIN-3 | 21629 | 71.8 |
| 176-cDNA-VR-B-CONMIN-2 | 48846 | 67.2 | 179-cDNA-VR-R-CONMIN-4 | 55639 | 67.6 |
| 178-cDNA-VR-B-CONMIN-3 | 64345 | 66.7 | 17-DNA-VR-R-CONMIN-1 | 14625 | 67.3 |
| 180-cDNA-VR-B-CONMIN-4 | 50898 | 65.7 | 181-cDNA-VR-R-CONMIN-5 | 37772 | 69.3 |
| 182-cDNA-VR-B-CONMIN-5 | 35870 | 72.5 | 183-cDNA-VR-R-CONMIN-6 | 29856 | 61.7 |
| 184-cDNA-VR-B-CONMIN-6 | 36038 | 67.8 | 185-cDNA-VR-R-CONSLU-1 | 61100 | 68 |
| 186-cDNA-VR-B-CONSLU-1 | 40245 | 63.9 | 187-cDNA-VR-R-CONSLU-2 | 43693 | 72.2 |

| | | | | | |
|---|---|---|---|---|---|
| 188-cDNA-VR-B-CONSLU-2 | 55193 | 64.9 | 189-cDNA-VR-R-CONSLU-3 | 43275 | 76.9 |
| 18-DNA-VR-B-CONMIN-1 | 3405 | 65.5 | 191-cDNA-VR-R-CONSLU-4 | 31002 | 71.9 |
| 190-cDNA-VR-B-CONSLU-3 | 35824 | 66.3 | 193-cDNA-VR-R-CONSLU-5 | 48046 | 63.2 |
| 192-cDNA-VR-B-CONSLU-4 | 57941 | 66.4 | 195-cDNA-VR-R-CONSLU-6 | 41780 | 66.9 |
| 194-cDNA-VR-B-CONSLU-5 | 36339 | 69.2 | 197-cDNA-VR-R-ORG-1 | 67345 | 67 |
| 196-cDNA-VR-B-CONSLU-6 | 59244 | 67.8 | 199-cDNA-VR-R-ORG-2 | 36832 | 66.5 |
| 198-cDNA-VR-B-ORG-1 | 48292 | 62.2 | 19-DNA-VR-R-CONMIN-2 | 3032 | 66.5 |
| 200-cDNA-VR-B-ORG-2 | 43280 | 65.4 | 201-cDNA-VR-R-ORG-3 | 8655 | 67.5 |
| 202-cDNA-VR-B-ORG-3 | 44229 | 69.8 | 203-cDNA-VR-R-ORG-4 | 31491 | 66.1 |
| 204-cDNA-VR-B-ORG-4 | 62720 | 64.6 | 205-cDNA-VR-R-ORG-5 | 66529 | 57.6 |
| 206-cDNA-VR-B-ORG-5 | 52710 | 62.7 | 207-cDNA-VR-R-ORG-6 | 27566 | 60.8 |
| 208-cDNA-VR-B-ORG-6 | 37262 | 59.2 | 21-DNA-VR-R-CONMIN-3 | 2402 | 68.4 |
| 20-DNA-VR-B-CONMIN-2 | 3338 | 70.7 | 23-DNA-VR-R-CONMIN-4 | 2629 | 67.7 |
| 22-DNA-VR-B-CONMIN-3 | 2547 | 66.1 | 25-DNA-VR-R-CONMIN-5 | 7009 | 68.5 |
| 24-DNA-VR-B-CONMIN-4 | 6973 | 64.9 | 27-DNA-VR-R-CONMIN-6 | 1187 | 69 |
| 26-DNA-VR-B-CONMIN-5 | 4150 | 66.2 | 31-DNA-VR-R-CONSLU-2 | 2744 | 67.1 |
| 28-DNA-VR-B-CONMIN-6 | 2090 | 68.2 | 33-DNA-VR-R-CONSLU-3 | 8836 | 68.1 |
| 30-DNA-VR-B-CONSLU-1 | 1221 | 67.3 | 35-DNA-VR-R-CONSLU-4 | 6718 | 65.2 |
| 32-DNA-VR-B-CONSLU-2 | 4584 | 66.2 | 37-DNA-VR-R-CONSLU-5 | 5906 | 68.5 |
| 34-DNA-VR-B-CONSLU-3 | 10920 | 65.8 | 39-DNA-VR-R-CONSLU-6 | 3191 | 67.8 |
| 36-DNA-VR-B-CONSLU-4 | 5645 | 64.9 | 41-DNA-VR-R-ORG-1 | 8348 | 66.2 |
| 38-DNA-VR-B-CONSLU-5 | 4227 | 66.7 | 43-DNA-VR-R-ORG-2 | 2664 | 67.6 |
| 40-DNA-VR-B-CONSLU-6 | 6441 | 68.6 | 45-DNA-VR-R-ORG-3 | 2657 | 65.4 |
| 42-DNA-VR-B-ORG-1 | 4043 | 64.8 | 47-DNA-VR-R-ORG-4 | 3761 | 67.4 |
| 44-DNA-VR-B-ORG-2 | 7717 | 66.3 | 49-DNA-VR-R-ORG-5 | 8610 | 66.2 |
| 46-DNA-VR-B-ORG-3 | 1977 | 65.4 | 51-DNA-VR-R-ORG-6 | 2491 | 69.5 |
| 48-DNA-VR-B-ORG-4 | 4619 | 65.9 | 69-DNA-VR-R-CONMIN-1 | 8098 | 68.6 |
| 50-DNA-VR-B-ORG-5 | 2067 | 68.3 | 71-DNA-VR-R-CONMIN-2 | 2613 | 68.2 |

| | | | | | | | |
|---|---|---|---|---|---|---|---|
| | 52-DNA-VR-B-ORG-6 | 3711 | 66.4 | | 73-DNA-VR-R-CONMIN-3 | 7511 | 68.5 |
| | 70-DNA-VR-B-CONMIN-1 | 9136 | 68.6 | | 75-DNA-VR-R-CONMIN-4 | 5908 | 69.5 |
| | 72-DNA-VR-B-CONMIN-2 | 7814 | 67.9 | | 77-DNA-VR-R-CONMIN-5 | 3936 | 69.9 |
| | 74-DNA-VR-B-CONMIN-3 | 6251 | 61.7 | | 79-DNA-VR-R-CONMIN-6 | 2624 | 68 |
| | 76-DNA-VR-B-CONMIN-4 | 9613 | 62.2 | | 81-DNA-VR-R-CONSLU-1 | 3471 | 72.6 |
| | 78-DNA-VR-B-CONMIN-5 | 3105 | 68.6 | | 83-DNA-VR-R-CONSLU-2 | 2255 | 71.4 |
| | 80-DNA-VR-B-CONMIN-6 | 6258 | 66.9 | | 85-DNA-VR-R-CONSLU-3 | 2176 | 66.6 |
| | 82-DNA-VR-B-CONSLU-1 | 5563 | 65.4 | | 87-DNA-VR-R-CONSLU-4 | 1939 | 70.4 |
| | 84-DNA-VR-B-CONSLU-2 | 3483 | 65.3 | | 89-DNA-VR-R-CONSLU-5 | 6883 | 70.1 |
| | 86-DNA-VR-B-CONSLU-3 | 1704 | 65.8 | | 91-DNA-VR-R-CONSLU-6 | 5533 | 74.2 |
| | 88-DNA-VR-B-CONSLU-4 | 3948 | 68.2 | | 93-DNA-VR-R-ORG-1 | 6638 | 68 |
| | 90-DNA-VR-B-CONSLU-5 | 7584 | 67.9 | | 95-DNA-VR-R-ORG-2 | 6913 | 65.9 |
| | 92-DNA-VR-B-CONSLU-6 | 4958 | 67.2 | | 97-DNA-VR-R-ORG-3 | 7903 | 69.3 |
| | 94-DNA-VR-B-ORG-1 | 3863 | 64.1 | | 99-DNA-VR-R-ORG-4 | 7913 | 68.3 |
| | 96-DNA-VR-B-ORG-2 | 5987 | 66.2 | | | | |
| | 98-DNA-VR-B-ORG-3 | 4044 | 65.3 | | | | |
| | Total | 1603445 | | | Total | 1355996 | |
| Fungi-Bulk | 100-DNA-VR-B-ORG-4 | 24169 | 95.9 | Fungi-Rhizosphere | 101-DNA-VR-R-ORG-5 | 19269 | 96.2 |
| | 102-DNA-VR-B-ORG-5 | 34022 | 95.6 | | 103-DNA-VR-R-ORG-6 | 28668 | 97.4 |
| | 104-DNA-VR-B-ORG-6 | 19374 | 96.6 | | 123-cDNA-VR-R-CONMIN-2 | 9749 | 97.3 |
| | 124-cDNA-VR-B-CONMIN-2 | 5699 | 95.6 | | 125-cDNA-VR-R-CONMIN-3 | 2213 | 96.5 |
| | 126-cDNA-VR-B-CONMIN-3 | 1653 | 96.3 | | 127-cDNA-VR-R-CONMIN-4 | 2344 | 96.3 |
| | 128-cDNA-VR-B-CONMIN-4 | 4197 | 96.2 | | 129-cDNA-VR-R-CONMIN-5 | 12085 | 96.8 |
| | 130-cDNA-VR-B-CONMIN-5 | 12585 | 96.2 | | 131-cDNA-VR-R-CONMIN-6 | 9993 | 95.7 |
| | 132-cDNA-VR-B-CONMIN-6 | 8368 | 95.4 | | 133-cDNA-VR-R-CONSLU-1 | 5264 | 96.7 |
| | 134-cDNA-VR-B-CONSLU-1 | 2348 | 96.1 | | 135-cDNA-VR-R-CONSLU-2 | 2452 | 96.4 |
| | 136-cDNA-VR-B-CONSLU-2 | 4955 | 95.7 | | 139-cDNA-VR-R-CONSLU-4 | 1836 | 96.6 |

| | | | | | |
|---|---|---|---|---|---|
| 138-cDNA-VR-B-CONSLU-3 | 8729 | 95.9 | 141-cDNA-VR-R-CONSLU-5 | 3742 | 95.4 |
| 140-cDNA-VR-B-CONSLU-4 | 5660 | 96.1 | 143-cDNA-VR-R-CONSLU-6 | 2667 | 96.3 |
| 142-cDNA-VR-B-CONSLU-5 | 1808 | 96.2 | 147-cDNA-VR-R-ORG-2 | 3753 | 95.9 |
| 144-cDNA-VR-B-CONSLU-6 | 3404 | 95.4 | 149-cDNA-VR-R-ORG-3 | 5344 | 95 |
| 146-cDNA-VR-B-ORG-1 | 4283 | 96.7 | 151-cDNA-VR-R-ORG-4 | 4031 | 96.1 |
| 148-cDNA-VR-B-ORG-2 | 7270 | 96.4 | 155-cDNA-VR-R-ORG-6 | 3509 | 95.9 |
| 152-cDNA-VR-B-ORG-4 | 4947 | 96 | 173-cDNA-VR-R-CONMIN-1 | 5271 | 97 |
| 154-cDNA-VR-B-ORG-5 | 7985 | 95.6 | 175-cDNA-VR-R-CONMIN-2 | 5337 | 97.8 |
| 156-cDNA-VR-B-ORG-6 | 3461 | 95.4 | 177-cDNA-VR-R-CONMIN-3 | 3168 | 96.9 |
| 174-cDNA-VR-B-CONMIN-1 | 9933 | 95.2 | 179-cDNA-VR-R-CONMIN-4 | 6896 | 96.6 |
| 176-cDNA-VR-B-CONMIN-2 | 3365 | 96.7 | 17-DNA-VR-R-CONMIN-1 | 1292 | 96.4 |
| 178-cDNA-VR-B-CONMIN-3 | 6341 | 95.7 | 181-cDNA-VR-R-CONMIN-5 | 5917 | 95.5 |
| 180-cDNA-VR-B-CONMIN-4 | 5757 | 96.7 | 183-cDNA-VR-R-CONMIN-6 | 2025 | 96.5 |
| 182-cDNA-VR-B-CONMIN-5 | 7294 | 96 | 185-cDNA-VR-R-CONSLU-1 | 9640 | 96.8 |
| 184-cDNA-VR-B-CONMIN-6 | 8777 | 96 | 187-cDNA-VR-R-CONSLU-2 | 10505 | 95.6 |
| 186-cDNA-VR-B-CONSLU-1 | 5437 | 94.4 | 189-cDNA-VR-R-CONSLU-3 | 3687 | 95.8 |
| 188-cDNA-VR-B-CONSLU-2 | 6113 | 96.3 | 191-cDNA-VR-R-CONSLU-4 | 3726 | 95.9 |
| 18-DNA-VR-B-CONMIN-1 | 1701 | 95.7 | 193-cDNA-VR-R-CONSLU-5 | 18829 | 95.6 |
| 190-cDNA-VR-B-CONSLU-3 | 4535 | 96.1 | 195-cDNA-VR-R-CONSLU-6 | 15523 | 95.6 |
| 192-cDNA-VR-B-CONSLU-4 | 9160 | 96.4 | 197-cDNA-VR-R-ORG-1 | 5723 | 96.8 |
| 194-cDNA-VR-B-CONSLU-5 | 7967 | 96 | 199-cDNA-VR-R-ORG-2 | 5296 | 94.8 |

| | | | | | |
|---|---|---|---|---|---|
| 196-cDNA-VR-B-CONSLU-6 | 6496 | 96.2 | 19-DNA-VR-R-CONMIN-2 | 2511 | 97.1 |
| 198-cDNA-VR-B-ORG-1 | 3708 | 95.6 | 201-cDNA-VR-R-ORG-3 | 5499 | 95.9 |
| 200-cDNA-VR-B-ORG-2 | 8687 | 97 | 203-cDNA-VR-R-ORG-4 | 27227 | 95.8 |
| 202-cDNA-VR-B-ORG-3 | 27325 | 96.1 | 205-cDNA-VR-R-ORG-5 | 25887 | 95.5 |
| 204-cDNA-VR-B-ORG-4 | 23529 | 95.4 | 207-cDNA-VR-R-ORG-6 | 8289 | 96.7 |
| 206-cDNA-VR-B-ORG-5 | 11710 | 96.7 | 21-DNA-VR-R-CONMIN-3 | 1416 | 96.2 |
| 208-cDNA-VR-B-ORG-6 | 9377 | 95.9 | 23-DNA-VR-R-CONMIN-4 | 3760 | 95.2 |
| 20-DNA-VR-B-CONMIN-2 | 2417 | 97.5 | 25-DNA-VR-R-CONMIN-5 | 1256 | 95.8 |
| 22-DNA-VR-B-CONMIN-3 | 2891 | 95.8 | 27-DNA-VR-R-CONMIN-6 | 1696 | 95.9 |
| 24-DNA-VR-B-CONMIN-4 | 2700 | 95.6 | 31-DNA-VR-R-CONSLU-2 | 2374 | 95.6 |
| 26-DNA-VR-B-CONMIN-5 | 1958 | 95.9 | 35-DNA-VR-R-CONSLU-4 | 1684 | 96.9 |
| 28-DNA-VR-B-CONMIN-6 | 1960 | 95.6 | 39-DNA-VR-R-CONSLU-6 | 2996 | 95.9 |
| 30-DNA-VR-B-CONSLU-1 | 3064 | 96.5 | 41-DNA-VR-R-ORG-1 | 1988 | 96.3 |
| 32-DNA-VR-B-CONSLU-2 | 2580 | 96.2 | 43-DNA-VR-R-ORG-2 | 2178 | 95.4 |
| 34-DNA-VR-B-CONSLU-3 | 1406 | 96.7 | 45-DNA-VR-R-ORG-3 | 1494 | 96.3 |
| 38-DNA-VR-B-CONSLU-5 | 1955 | 96.3 | 47-DNA-VR-R-ORG-4 | 1174 | 95 |
| 40-DNA-VR-B-CONSLU-6 | 2352 | 96.2 | 49-DNA-VR-R-ORG-5 | 2869 | 96.3 |
| 42-DNA-VR-B-ORG-1 | 2012 | 96.7 | 51-DNA-VR-R-ORG-6 | 1388 | 95.5 |
| 44-DNA-VR-B-ORG-2 | 2099 | 96 | 69-DNA-VR-R-CONMIN-1 | 2301 | 94.7 |
| 46-DNA-VR-B-ORG-3 | 1493 | 96.7 | 73-DNA-VR-R-CONMIN-3 | 1577 | 95.8 |
| 48-DNA-VR-B-ORG-4 | 4348 | 95.8 | 77-DNA-VR-R-CONMIN-5 | 2938 | 95.7 |
| 50-DNA-VR-B-ORG-5 | 1030 | 96.7 | 79-DNA-VR-R-CONMIN-6 | 1109 | 96.5 |
| 52-DNA-VR-B-ORG-6 | 1288 | 95.3 | 81-DNA-VR-R-CONSLU-1 | 3103 | 97.1 |
| 70-DNA-VR-B-CONMIN-1 | 1405 | 96.2 | 83-DNA-VR-R-CONSLU-2 | 1548 | 96.3 |
| 76-DNA-VR-B-CONMIN-4 | 1015 | 96.5 | 85-DNA-VR-R-CONSLU-3 | 2293 | 96 |
| 78-DNA-VR-B-CONMIN-5 | 1074 | 95.7 | 87-DNA-VR-R-CONSLU-4 | 1290 | 96 |
| 80-DNA-VR-B-CONMIN-6 | 1599 | 95.7 | 89-DNA-VR-R-CONSLU-5 | 2538 | 97.3 |
| 82-DNA-VR-B-CONSLU-1 | 1544 | 96.7 | 91-DNA-VR-R-CONSLU-6 | 1907 | 95.9 |
| 84-DNA-VR-B-CONSLU-2 | 1127 | 97.1 | 93-DNA-VR-R-ORG-1 | 2345 | 96.6 |
| 86-DNA-VR-B-CONSLU-3 | 1312 | 95.4 | 95-DNA-VR-R-ORG-2 | 4548 | 95.5 |

| | | | | | | | |
|---|---|---|---|---|---|---|---|
| | 88-DNA-VR-B-CONSLU-4 | 1522 | 95.9 | | 97-DNA-VR-R-ORG-3 | 15457 | 96 |
| | 90-DNA-VR-B-CONSLU-5 | 3189 | 95.7 | | 99-DNA-VR-R-ORG-4 | 20982 | 95.2 |
| | 92-DNA-VR-B-CONSLU-6 | 3787 | 95.4 | | | | |
| | 94-DNA-VR-B-ORG-1 | 1617 | 95.2 | | | | |
| | 96-DNA-VR-B-ORG-2 | 1628 | 96.4 | | | | |
| | 98-DNA-VR-B-ORG-3 | 7528 | 95.7 | | | | |
| | Total | 396029 | | | Total | 379376 | |
| Metazoa-bulk | 100-DNA-VR-B-ORG-4 | 30746 | 58.3 | Metazoa-rhizosphere | 101-DNA-VR-R-ORG-5 | 5190 | 97.5 |
| | 102-DNA-VR-B-ORG-5 | 29230 | 58.8 | | 103-DNA-VR-R-ORG-6 | 11244 | 96.7 |
| | 104-DNA-VR-B-ORG-6 | 13012 | 60.8 | | 121-cDNA-VR-R-CONMIN-1 | 1602 | 97.7 |
| | 124-cDNA-VR-B-CONMIN-2 | 12887 | 60.7 | | 123-cDNA-VR-R-CONMIN-2 | 26828 | 97.9 |
| | 126-cDNA-VR-B-CONMIN-3 | 1907 | 61.8 | | 125-cDNA-VR-R-CONMIN-3 | 6763 | 97.3 |
| | 128-cDNA-VR-B-CONMIN-4 | 13366 | 61.6 | | 127-cDNA-VR-R-CONMIN-4 | 24506 | 96.8 |
| | 130-cDNA-VR-B-CONMIN-5 | 8500 | 62.5 | | 129-cDNA-VR-R-CONMIN-5 | 31022 | 97.5 |
| | 132-cDNA-VR-B-CONMIN-6 | 16317 | 60.7 | | 131-cDNA-VR-R-CONMIN-6 | 26103 | 96.3 |
| | 134-cDNA-VR-B-CONSLU-1 | 4549 | 63 | | 133-cDNA-VR-R-CONSLU-1 | 24629 | 98.1 |
| | 136-cDNA-VR-B-CONSLU-2 | 16335 | 60.2 | | 135-cDNA-VR-R-CONSLU-2 | 30950 | 97 |
| | 138-cDNA-VR-B-CONSLU-3 | 13386 | 64.5 | | 137-cDNA-VR-R-CONSLU-3 | 1209 | 96.7 |
| | 140-cDNA-VR-B-CONSLU-4 | 12883 | 62.6 | | 139-cDNA-VR-R-CONSLU-4 | 19938 | 97.8 |
| | 142-cDNA-VR-B-CONSLU-5 | 8974 | 64.2 | | 141-cDNA-VR-R-CONSLU-5 | 16996 | 98.3 |
| | 144-cDNA-VR-B-CONSLU-6 | 8904 | 64.1 | | 143-cDNA-VR-R-CONSLU-6 | 12378 | 97.4 |
| | 146-cDNA-VR-B-ORG-1 | 13979 | 58.8 | | 145-cDNA-VR-R-ORG-1 | 2296 | 96.2 |
| | 148-cDNA-VR-B-ORG-2 | 18130 | 58.5 | | 147-cDNA-VR-R-ORG-2 | 40954 | 97.4 |
| | 152-cDNA-VR-B-ORG-4 | 22000 | 58.9 | | 149-cDNA-VR-R-ORG-3 | 26411 | 97.3 |
| | 154-cDNA-VR-B-ORG-5 | 5215 | 60.7 | | 151-cDNA-VR-R-ORG-4 | 28138 | 96.9 |

| | | | | | |
|---|---|---|---|---|---|
| 156-cDNA-VR-B-ORG-6 | 7793 | 60.6 | 153-cDNA-VR-R-ORG-5 | 7171 | 97.6 |
| 174-cDNA-VR-B-CONMIN-1 | 8289 | 60.1 | 155-cDNA-VR-R-ORG-6 | 44763 | 97.4 |
| 176-cDNA-VR-B-CONMIN-2 | 13260 | 59.6 | 173-cDNA-VR-R-CONMIN-1 | 16870 | 96.1 |
| 178-cDNA-VR-B-CONMIN-3 | 25456 | 57.7 | 175-cDNA-VR-R-CONMIN-2 | 19424 | 97.1 |
| 180-cDNA-VR-B-CONMIN-4 | 11385 | 57.9 | 177-cDNA-VR-R-CONMIN-3 | 14882 | 96.4 |
| 182-cDNA-VR-B-CONMIN-5 | 26428 | 63.1 | 179-cDNA-VR-R-CONMIN-4 | 43887 | 97.9 |
| 184-cDNA-VR-B-CONMIN-6 | 21589 | 59.7 | 17-DNA-VR-R-CONMIN-1 | 2327 | 96.7 |
| 186-cDNA-VR-B-CONSLU-1 | 7733 | 61.9 | 181-cDNA-VR-R-CONMIN-5 | 32311 | 96.5 |
| 188-cDNA-VR-B-CONSLU-2 | 10043 | 62.9 | 183-cDNA-VR-R-CONMIN-6 | 36552 | 97.4 |
| 18-DNA-VR-B-CONMIN-1 | 2730 | 61.9 | 185-cDNA-VR-R-CONSLU-1 | 22204 | 97.2 |
| 190-cDNA-VR-B-CONSLU-3 | 12394 | 63.2 | 187-cDNA-VR-R-CONSLU-2 | 22421 | 96.9 |
| 192-cDNA-VR-B-CONSLU-4 | 14812 | 58.6 | 189-cDNA-VR-R-CONSLU-3 | 11110 | 97.3 |
| 194-cDNA-VR-B-CONSLU-5 | 16501 | 61.3 | 191-cDNA-VR-R-CONSLU-4 | 10001 | 97 |
| 196-cDNA-VR-B-CONSLU-6 | 14568 | 62.5 | 193-cDNA-VR-R-CONSLU-5 | 23333 | 97.2 |
| 198-cDNA-VR-B-ORG-1 | 14985 | 58.1 | 195-cDNA-VR-R-CONSLU-6 | 13697 | 96.6 |
| 200-cDNA-VR-B-ORG-2 | 10968 | 57.4 | 197-cDNA-VR-R-ORG-1 | 32778 | 95.8 |
| 202-cDNA-VR-B-ORG-3 | 23100 | 57.1 | 199-cDNA-VR-R-ORG-2 | 22876 | 96.3 |
| 204-cDNA-VR-B-ORG-4 | 36170 | 63.2 | 201-cDNA-VR-R-ORG-3 | 7321 | 97 |
| 206-cDNA-VR-B-ORG-5 | 37305 | 59.9 | 203-cDNA-VR-R-ORG-4 | 22590 | 95.9 |
| 208-cDNA-VR-B-ORG-6 | 24941 | 57.3 | 205-cDNA-VR-R-ORG-5 | 15113 | 97.1 |
| 20-DNA-VR-B-CONMIN-2 | 1505 | 66.5 | 207-cDNA-VR-R-ORG-6 | 12231 | 96.6 |
| 26-DNA-VR-B-CONMIN-5 | 2168 | 61 | 21-DNA-VR-R-CONMIN-3 | 1109 | 97.2 |
| 28-DNA-VR-B-CONMIN-6 | 1373 | 65.9 | 23-DNA-VR-R-CONMIN-4 | 1802 | 97.7 |
| 30-DNA-VR-B-CONSLU-1 | 1205 | 62 | 25-DNA-VR-R-CONMIN-5 | 4217 | 96.9 |

| | | | | | | | |
|---|---|---|---|---|---|---|---|
| | 34-DNA-VR-B-CONSLU-3 | 1041 | 61.9 | | 29-DNA-VR-R-CONSLU-1 | 1015 | 97.3 |
| | 36-DNA-VR-B-CONSLU-4 | 1107 | 62.9 | | 31-DNA-VR-R-CONSLU-2 | 1070 | 96.7 |
| | 38-DNA-VR-B-CONSLU-5 | 1234 | 65.5 | | 33-DNA-VR-R-CONSLU-3 | 1454 | 97.1 |
| | 42-DNA-VR-B-ORG-1 | 2552 | 61.4 | | 37-DNA-VR-R-CONSLU-5 | 1213 | 97.6 |
| | 44-DNA-VR-B-ORG-2 | 8051 | 60 | | 41-DNA-VR-R-ORG-1 | 2356 | 96.7 |
| | 46-DNA-VR-B-ORG-3 | 1160 | 60.1 | | 43-DNA-VR-R-ORG-2 | 2248 | 96.6 |
| | 48-DNA-VR-B-ORG-4 | 1016 | 61.8 | | 45-DNA-VR-R-ORG-3 | 1338 | 97.2 |
| | 50-DNA-VR-B-ORG-5 | 1077 | 64 | | 47-DNA-VR-R-ORG-4 | 1196 | 97.4 |
| | 52-DNA-VR-B-ORG-6 | 1838 | 62.3 | | 49-DNA-VR-R-ORG-5 | 10027 | 97.3 |
| | 70-DNA-VR-B-CONMIN-1 | 1372 | 62.5 | | 51-DNA-VR-R-ORG-6 | 8086 | 98.1 |
| | 74-DNA-VR-B-CONMIN-3 | 1013 | 60.8 | | 69-DNA-VR-R-CONMIN-1 | 1115 | 97.6 |
| | 76-DNA-VR-B-CONMIN-4 | 2083 | 59.8 | | 73-DNA-VR-R-CONMIN-3 | 5769 | 97.1 |
| | 78-DNA-VR-B-CONMIN-5 | 1402 | 64.2 | | 75-DNA-VR-R-CONMIN-4 | 1013 | 97.8 |
| | 80-DNA-VR-B-CONMIN-6 | 1435 | 61.4 | | 77-DNA-VR-R-CONMIN-5 | 1497 | 97.2 |
| | 82-DNA-VR-B-CONSLU-1 | 1514 | 59.2 | | 81-DNA-VR-R-CONSLU-1 | 1308 | 96.2 |
| | 84-DNA-VR-B-CONSLU-2 | 2360 | 61.3 | | 83-DNA-VR-R-CONSLU-2 | 1940 | 97.5 |
| | 88-DNA-VR-B-CONSLU-4 | 1730 | 62.2 | | 85-DNA-VR-R-CONSLU-3 | 1150 | 97.5 |
| | 90-DNA-VR-B-CONSLU-5 | 5499 | 60.6 | | 89-DNA-VR-R-CONSLU-5 | 4873 | 97.7 |
| | 92-DNA-VR-B-CONSLU-6 | 1800 | 61.1 | | 91-DNA-VR-R-CONSLU-6 | 1486 | 97.2 |
| | 94-DNA-VR-B-ORG-1 | 1410 | 56.2 | | 93-DNA-VR-R-ORG-1 | 3851 | 96.9 |
| | 96-DNA-VR-B-ORG-2 | 1624 | 62.8 | | 95-DNA-VR-R-ORG-2 | 3089 | 97.5 |
| | 98-DNA-VR-B-ORG-3 | 6118 | 63 | | 97-DNA-VR-R-ORG-3 | 7373 | 97.2 |
| | | | | | 99-DNA-VR-R-ORG-4 | 14740 | 96.8 |
| | Total | 645457 | | | Total | 857354 | |
| | Valthermond | | | | Valthermond | | |
| | Sample | Number sequences - singletons | Good's coverage (%) | | Sample | Number sequences - singletons | Good's coverage (%) |
| Bacteria-Bulk | 106-cDNA-VA-B-COMP-1 | 6474 | 63 | Bacteria-Rhizosphere | 113-cDNA-VA-R-COMP-3 | 1318 | 67 |
| | 108-cDNA-VA-B-COMP-2 | 6217 | 64.2 | | 115-cDNA-VA-R-COMP-4 | 6598 | 66.1 |
| | 10-DNA-VA-B-COMP-3 | 1169 | 71.3 | | 117-cDNA-VA-R-NO-COMP-3 | 13620 | 63.1 |

| | | | | | |
|---|---|---|---|---|---|
| 112-cDNA-VA-B-NO-COMP-2 | 7398 | 72.8 | 119-cDNA-VA-R-NO-COMP-4 | 10914 | 64.2 |
| 114-cDNA-VA-B-COMP-3 | 11228 | 66.1 | 11-DNA-VA-R-COMP-4 | 1264 | 70.4 |
| 120-cDNA-VA-B-NO-COMP-4 | 7219 | 66.7 | 13-DNA-VA-R-NO-COMP-3 | 2125 | 66.9 |
| 12-DNA-VA-B-COMP-4 | 1097 | 69.6 | 157-cDNA-VA-R-COMP-1 | 7771 | 70 |
| 14-DNA-VA-B-NO-COMP-3 | 1617 | 71.1 | 159-cDNA-VA-R-COMP-2 | 9839 | 67.7 |
| 158-cDNA-VA-B-COMP-1 | 16189 | 64.6 | 163-cDNA-VA-R-NO-COMP-2 | 5315 | 64.9 |
| 160-cDNA-VA-B-COMP-2 | 8872 | 66.6 | 165-cDNA-VA-R-COMP-3 | 15419 | 64.3 |
| 162-cDNA-VA-B-NO-COMP-1 | 6973 | 67.5 | 169-cDNA-VA-R-NO-COMP-3 | 5617 | 73 |
| 164-cDNA-VA-B-NO-COMP-2 | 2400 | 69 | 171-cDNA-VA-R-NO-COMP-4 | 5209 | 69.3 |
| 166-cDNA-VA-B-COMP-3 | 2721 | 64.3 | 1-DNA-VA-R-COMP-1 | 3419 | 65.7 |
| 168-cDNA-VA-B-COMP-4 | 3910 | 68.5 | 3-DNA-VA-R-COMP-2 | 1494 | 67.2 |
| 170-cDNA-VA-B-NO-COMP-3 | 14496 | 69.6 | 55-DNA-VA-R-COMP-2 | 1392 | 69.4 |
| 172-cDNA-VA-B-NO-COMP-4 | 7126 | 67 | 59-DNA-VA-R-NO-COMP-2 | 1670 | 69 |
| 2-DNA-VA-B-COMP-1 | 1598 | 69.8 | 65-DNA-VA-R-NO-COMP-3 | 1858 | 73.1 |
| 4-DNA-VA-B-COMP-2 | 3546 | 66.8 | 67-DNA-VA-R-NO-COMP-4 | 2570 | 72.2 |
| 54-DNA-VA-B-COMP-1 | 1152 | 71.4 | 7-DNA-VA-R-NO-COMP-2 | 1065 | 67.9 |
| 58-DNA-VA-B-NO-COMP-1 | 1573 | 70.1 | | | |
| 60-DNA-VA-B-NO-COMP-2 | 3155 | 72.9 | | | |
| 62-DNA-VA-B-COMP-3 | 1299 | 70.9 | | | |
| 66-DNA-VA-B-NO-COMP-3 | 1506 | 70.5 | | | |
| 68-DNA-VA-B-NO-COMP-4 | 1314 | 72.7 | | | |

| | | | | | | | |
|---|---|---|---|---|---|---|---|
| | 8-DNA-VA-B-NO-COMP-2 | 1071 | 71.7 | | | | |
| | Total | 121320 | | | Total | 98477 | |
| Protozoa-bulk | 106-cDNA-VA-B-COMP-1 | 19557 | 68.7 | Protozoa-rhizosphere | 105-cDNA-VA-R-COMP-1 | 2270 | 78.4 |
| | 108-cDNA-VA-B-COMP-2 | 35341 | 76.9 | | 107-cDNA-VA-R-COMP-2 | 4427 | 84.2 |
| | 10-DNA-VA-B-COMP-3 | 11223 | 72.6 | | 113-cDNA-VA-R-COMP-3 | 6140 | 76.1 |
| | 112-cDNA-VA-B-NO-COMP-2 | 20493 | 83.5 | | 115-cDNA-VA-R-COMP-4 | 18058 | 74.8 |
| | 114-cDNA-VA-B-COMP-3 | 30588 | 77.1 | | 117-cDNA-VA-R-NO-COMP-3 | 14993 | 74.1 |
| | 116-cDNA-VA-B-COMP-4 | 2966 | 70.2 | | 119-cDNA-VA-R-NO-COMP-4 | 27834 | 81.2 |
| | 120-cDNA-VA-B-NO-COMP-4 | 14092 | 72.9 | | 11-DNA-VA-R-COMP-4 | 14215 | 74.2 |
| | 12-DNA-VA-B-COMP-4 | 12092 | 70.9 | | 13-DNA-VA-R-NO-COMP-3 | 8387 | 75.3 |
| | 14-DNA-VA-B-NO-COMP-3 | 3204 | 66.5 | | 157-cDNA-VA-R-COMP-1 | 44453 | 89.8 |
| | 158-cDNA-VA-B-COMP-1 | 33777 | 74.4 | | 159-cDNA-VA-R-COMP-2 | 37016 | 87.2 |
| | 160-cDNA-VA-B-COMP-2 | 36186 | 87.6 | | 15-DNA-VA-R-NO-COMP-4 | 6403 | 75.5 |
| | 162-cDNA-VA-B-NO-COMP-1 | 36675 | 84.1 | | 161-cDNA-VA-R-NO-COMP-1 | 9173 | 92.6 |
| | 164-cDNA-VA-B-NO-COMP-2 | 40728 | 79.1 | | 163-cDNA-VA-R-NO-COMP-2 | 41662 | 91.1 |
| | 166-cDNA-VA-B-COMP-3 | 11924 | 77.2 | | 165-cDNA-VA-R-COMP-3 | 37310 | 78.5 |
| | 168-cDNA-VA-B-COMP-4 | 33923 | 83.9 | | 167-cDNA-VA-R-COMP-4 | 4447 | 86.1 |
| | 16-DNA-VA-B-NO-COMP-4 | 15874 | 71.3 | | 169-cDNA-VA-R-NO-COMP-3 | 13567 | 83.4 |
| | 170-cDNA-VA-B-NO-COMP-3 | 32400 | 84.3 | | 171-cDNA-VA-R-NO-COMP-4 | 77628 | 92 |
| | 172-cDNA-VA-B-NO-COMP-4 | 40644 | 84.7 | | 1-DNA-VA-R-COMP-1 | 19332 | 75.9 |
| | 2-DNA-VA-B-COMP-1 | 21217 | 69.7 | | 3-DNA-VA-R-COMP-2 | 17431 | 75.8 |
| | 4-DNA-VA-B-COMP-2 | 21122 | 72.2 | | 53-DNA-VA-R-COMP-1 | 1847 | 79.2 |

| | | | | | | |
|---|---|---|---|---|---|---|
| | 54-DNA-VA-B-COMP-1 | 1270 | 72.5 | | 55-DNA-VA-R-COMP-2 | 3252 | 76.5 |
| | 56-DNA-VA-B-COMP-2 | 9067 | 76.0 | | 57-DNA-VA-R-NO-COMP-1 | 4515 | 81.4 |
| | 58-DNA-VA-B-NO-COMP-1 | 2863 | 71.8 | | 59-DNA-VA-R-NO-COMP-2 | 4381 | 76.6 |
| | 60-DNA-VA-B-NO-COMP-2 | 4356 | 73.4 | | 5-DNA-VA-R-NO-COMP-1 | 13918 | 78.4 |
| | 62-DNA-VA-B-COMP-3 | 2579 | 72.4 | | 61-DNA-VA-R-COMP-3 | 3606 | 77.5 |
| | 64-DNA-VA-B-COMP-4 | 7945 | 70.1 | | 63-DNA-VA-R-COMP-4 | 6859 | 78.4 |
| | 66-DNA-VA-B-NO-COMP-3 | 4041 | 71.4 | | 65-DNA-VA-R-NO-COMP-3 | 6226 | 76.9 |
| | 68-DNA-VA-B-NO-COMP-4 | 8565 | 73.1 | | 67-DNA-VA-R-NO-COMP-4 | 7482 | 76.8 |
| | 6-DNA-VA-B-NO-COMP-1 | 9542 | 69.4 | | 7-DNA-VA-R-NO-COMP-2 | 11212 | 76.7 |
| | 8-DNA-VA-B-NO-COMP-2 | 21775 | 72.1 | | 9-DNA-VA-R-COMP-3 | 15594 | 75.1 |
| | Total | 546029 | | | Total | 483638 | |
| Fungi-bulk | 106-cDNA-VA-B-COMP-1 | 3874 | 97 | Fungi-rhizosphere | 113-cDNA-VA-R-COMP-3 | 1012 | 96.5 |
| | 108-cDNA-VA-B-COMP-2 | 2744 | 97.2 | | 115-cDNA-VA-R-COMP-4 | 2442 | 96.2 |
| | 10-DNA-VA-B-COMP-3 | 5084 | 97.2 | | 117-cDNA-VA-R-NO-COMP-3 | 2118 | 95.6 |
| | 112-cDNA-VA-B-NO-COMP-2 | 3313 | 98.3 | | 119-cDNA-VA-R-NO-COMP-4 | 1581 | 96.1 |
| | 114-cDNA-VA-B-COMP-3 | 2487 | 95.7 | | 13-DNA-VA-R-NO-COMP-3 | 1250 | 96.6 |
| | 120-cDNA-VA-B-NO-COMP-4 | 3287 | 96.7 | | 157-cDNA-VA-R-COMP-1 | 1360 | 97 |
| | 12-DNA-VA-B-COMP-4 | 1734 | 97 | | 15-DNA-VA-R-NO-COMP-4 | 2432 | 96 |
| | 14-DNA-VA-B-NO-COMP-3 | 2932 | 96.6 | | 165-cDNA-VA-R-COMP-3 | 2679 | 96.7 |
| | 158-cDNA-VA-B-COMP-1 | 7164 | 96.3 | | 169-cDNA-VA-R-NO-COMP-3 | 2670 | 95.8 |
| | 160-cDNA-VA-B-COMP-2 | 4514 | 98.6 | | 171-cDNA-VA-R-NO-COMP-4 | 3342 | 96.1 |
| | 162-cDNA-VA-B-NO-COMP-1 | 3147 | 98.6 | | 1-DNA-VA-R-COMP-1 | 4166 | 96.3 |

| | | | | | | | |
|---|---|---|---|---|---|---|---|
| | 164-cDNA-VA-B-NO-COMP-2 | 2411 | 96.3 | | 3-DNA-VA-R-COMP-2 | 8228 | 95.7 |
| | 166-cDNA-VA-B-COMP-3 | 1606 | 97.2 | | 55-DNA-VA-R-COMP-2 | 1710 | 96 |
| | 16-DNA-VA-B-NO-COMP-4 | 1035 | 96.3 | | 57-DNA-VA-R-NO-COMP-1 | 1256 | 96.3 |
| | 170-cDNA-VA-B-NO-COMP-3 | 4474 | 97 | | 59-DNA-VA-R-NO-COMP-2 | 1994 | 97 |
| | 172-cDNA-VA-B-NO-COMP-4 | 1452 | 97.3 | | 5-DNA-VA-R-NO-COMP-1 | 6315 | 97.2 |
| | 2-DNA-VA-B-COMP-1 | 1529 | 96.7 | | 61-DNA-VA-R-COMP-3 | 1128 | 96.4 |
| | 4-DNA-VA-B-COMP-2 | 2227 | 97.5 | | 63-DNA-VA-R-COMP-4 | 1538 | 96 |
| | 54-DNA-VA-B-COMP-1 | 1074 | 95.7 | | 65-DNA-VA-R-NO-COMP-3 | 1952 | 96.5 |
| | 56-DNA-VA-B-COMP-2 | 3282 | 96.7 | | 67-DNA-VA-R-NO-COMP-4 | 1089 | 95.6 |
| | 60-DNA-VA-B-NO-COMP-2 | 1914 | 96.8 | | 7-DNA-VA-R-NO-COMP-2 | 8214 | 96.3 |
| | 62-DNA-VA-B-COMP-3 | 1810 | 96.9 | | | | |
| | 64-DNA-VA-B-COMP-4 | 1119 | 96.6 | | | | |
| | 68-DNA-VA-B-NO-COMP-4 | 2005 | 96.5 | | | | |
| | 6-DNA-VA-B-NO-COMP-1 | 3031 | 96.4 | | | | |
| | 8-DNA-VA-B-NO-COMP-2 | 3825 | 96.6 | | | | |
| | Total | 73074 | | | Total | 58476 | |
| Metazoa-bulk | 106-cDNA-VA-B-COMP-1 | 3956 | 97.7 | Metazoa-rhizosphere | 107-cDNA-VA-R-COMP-2 | 2508 | 97.2 |
| | 108-cDNA-VA-B-COMP-2 | 7561 | 97.7 | | 113-cDNA-VA-R-COMP-3 | 5008 | 96.6 |
| | 112-cDNA-VA-B-NO-COMP-2 | 6499 | 97.5 | | 115-cDNA-VA-R-COMP-4 | 11740 | 96.5 |
| | 114-cDNA-VA-B-COMP-3 | 18006 | 96.9 | | 117-cDNA-VA-R-NO-COMP-3 | 16449 | 97.5 |
| | 116-cDNA-VA-B-COMP-4 | 2322 | 96.3 | | 119-cDNA-VA-R-NO-COMP-4 | 28055 | 97.3 |
| | 120-cDNA-VA-B-NO-COMP-4 | 12610 | 97.6 | | 13-DNA-VA-R-NO-COMP-3 | 1504 | 97 |
| | 14-DNA-VA-B-NO-COMP-3 | 1417 | 97 | | 157-cDNA-VA-R-COMP-1 | 25782 | 96.2 |

| | | | | | |
|---|---|---|---|---|---|
| 158-cDNA-VA-B-COMP-1 | 20867 | 96.9 | 159-cDNA-VA-R-COMP-2 | 12324 | 97.2 |
| 160-cDNA-VA-B-COMP-2 | 11834 | 97.8 | 163-cDNA-VA-R-NO-COMP-2 | 25450 | 97.3 |
| 162-cDNA-VA-B-NO-COMP-1 | 12167 | 98 | 165-cDNA-VA-R-COMP-3 | 14226 | 96.3 |
| 164-cDNA-VA-B-NO-COMP-2 | 9643 | 97.3 | 167-cDNA-VA-R-COMP-4 | 1203 | 97.3 |
| 166-cDNA-VA-B-COMP-3 | 6134 | 98.1 | 169-cDNA-VA-R-NO-COMP-3 | 8473 | 97.6 |
| 168-cDNA-VA-B-COMP-4 | 10503 | 97 | 171-cDNA-VA-R-NO-COMP-4 | 20878 | 96.7 |
| 170-cDNA-VA-B-NO-COMP-3 | 15446 | 97.9 | 1-DNA-VA-R-COMP-1 | 5100 | 96.2 |
| 172-cDNA-VA-B-NO-COMP-4 | 10152 | 97.2 | 3-DNA-VA-R-COMP-2 | 3469 | 96.9 |
| 2-DNA-VA-B-COMP-1 | 2540 | 97.7 | 53-DNA-VA-R-COMP-1 | 2862 | 96.9 |
| 4-DNA-VA-B-COMP-2 | 4973 | 97.7 | 55-DNA-VA-R-COMP-2 | 2834 | 96.3 |
| 54-DNA-VA-B-COMP-1 | 1239 | 97.4 | 5-DNA-VA-R-NO-COMP-1 | 3396 | 97.3 |
| 56-DNA-VA-B-COMP-2 | 5296 | 98.1 | 61-DNA-VA-R-COMP-3 | 1330 | 96.7 |
| 6-DNA-VA-B-NO-COMP-1 | 7441 | 98.1 | 65-DNA-VA-R-NO-COMP-3 | 1767 | 97.5 |
| 8-DNA-VA-B-NO-COMP-2 | 4953 | 97.5 | 67-DNA-VA-R-NO-COMP-4 | 1061 | 96.6 |
| | | | 7-DNA-VA-R-NO-COMP-2 | 6967 | 97.4 |
| | | | 9-DNA-VA-R-COMP-3 | 1331 | 97.5 |
| Total | 175559 | | Total | 203717 | |

**S4**: Results of PERMANOVA testing samples of Valthermond for the effects of nucleic acid (cDNA and DNA) fertilization treatment (compost and no compost), sample type (bulk soil and rhizosphere) and time point (vegetative and generative) on soil microbial community.

| Source | Df | SS | F | R2 | P |
|---|---|---|---|---|---|
| **Bacteria** | | | | | |
| Nucleic Acid | 1 | 0.16466 | 34.089 | 0.43281 | **9.999e-05** |
| Sample Type | 1 | 0.01011 | 2.094 | 0.02659 | 0.09769 . |
| Treatment | 1 | 0.00303 | 0.627 | 0.00796 | 0.56624 |
| Time Point | 1 | 0.00871 | 1.803 | 0.02289 | 0.13189 |
| Residuals | 28 | 0.13524 | | 0.3555 | |
| **Fungi** | | | | | |
| Nucleic_Acid | 1 | 0.12438 | 8.1517 | 0.13886 | **1.00E-04** |
| Sample_Type | 1 | 0.01564 | 1.025 | 0.01746 | **0.006099** |
| Treatment | 1 | 0.04605 | 3.018 | 0.05141 | 0.374263 |
| Time_Point | 1 | 0.12438 | 8.1517 | 0.13886 | **0.009499** |
| Residuals | 31 | 0.47301 | | 0.52808 | |
| **Protozoa** | | | | | |
| Nucleic_Acid | 1 | 0.3846 | 27.6554 | 0.27143 | **1.00E-04** |
| Sample_Type | 1 | 0.034 | 2.445 | 0.024 | **0.05859** |
| Treatment | 1 | 0.01579 | 1.1356 | 0.01115 | 0.27947 |
| Time_Point | 1 | 0.05514 | 3.9648 | 0.03891 | **0.0121** |
| Residuals | 44 | 0.61191 | | 0.43185 | |
| **Metazoa** | | | | | |
| Nucleic_Acid | 1 | 0.38031 | 13.3288 | 0.21868 | **1.00E-04** |
| Sample_Type | 1 | 0.15642 | 5.482 | 0.08994 | **1.00E-04** |
| Treatment | 1 | 0.04597 | 1.6111 | 0.02643 | 0.10279 |
| Time_Point | 1 | 0.06569 | 2.3022 | 0.03777 | **0.026** |
| Residuals | 29 | 0.82746 | | 0.47579 | |
