## Supplementary figures 1-5 for "Mapping of long-term impact of conventional and organic soil management on resident and active fractions of rhizosphere communities of barley"

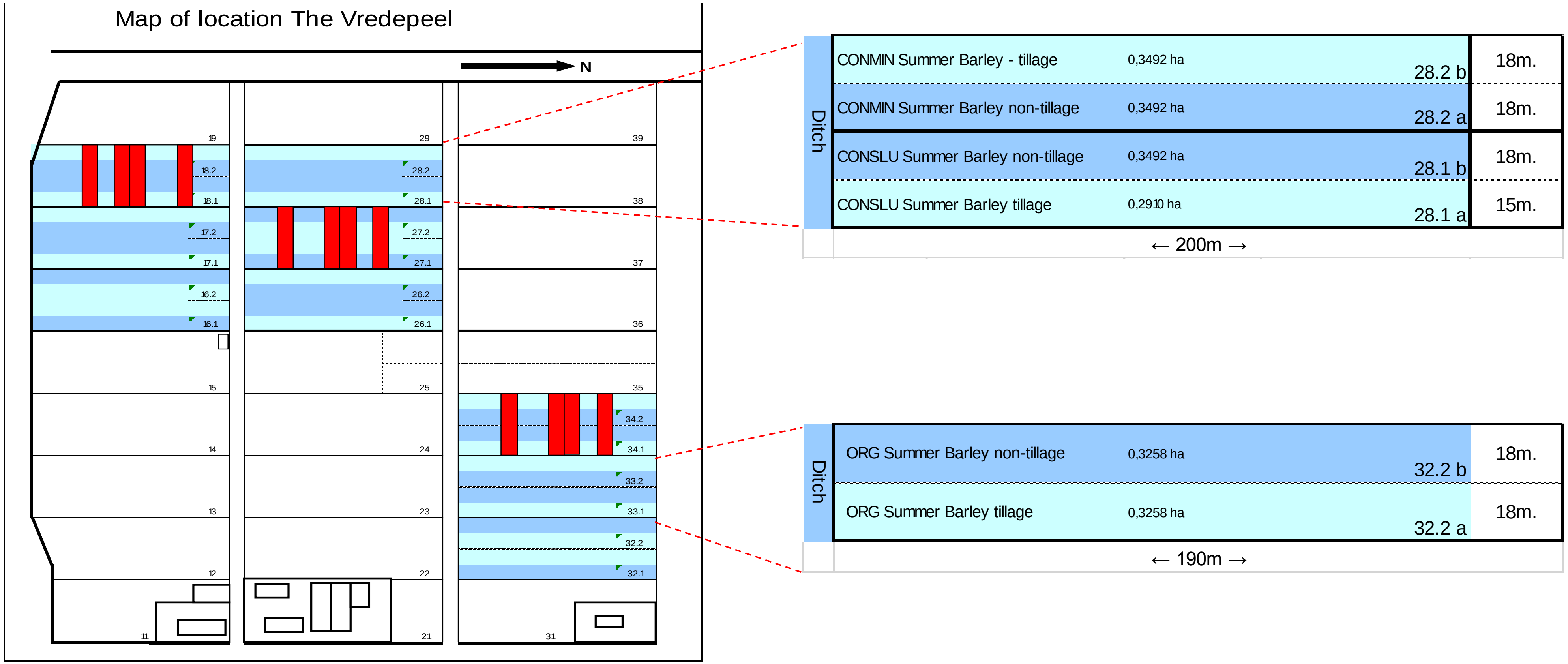


**Figure S1**: field set up location Vredepeel (sandy soils)

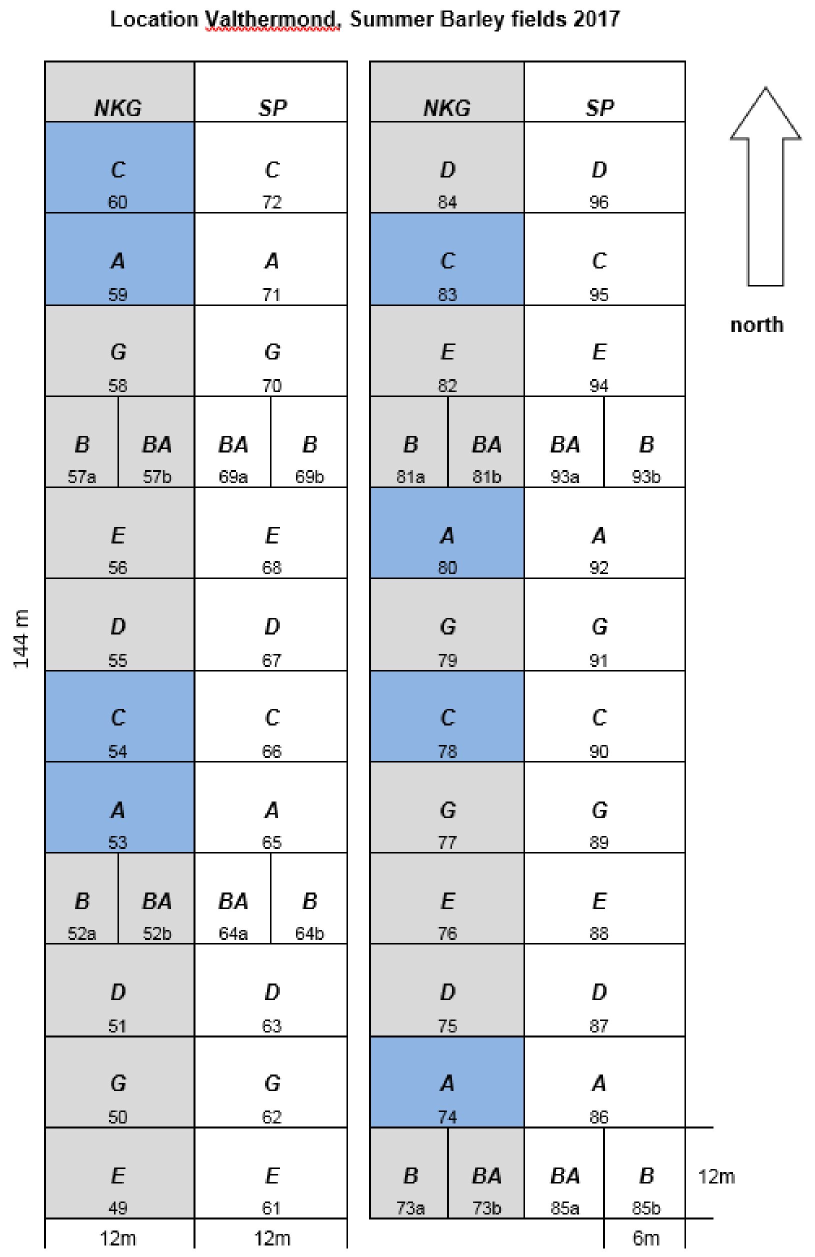


| | Object |
|---|---|
| *A* | Standard |
| *B* | Tagetes |
| *C* | Compost |
| *D* | Hortinova bodembalance |
| *E* | Combination (BCDG - Targetes 1x / 8 years) |
| *G* | Rock flour |
| *BA* | Standaard (tagetes 1x / 8 years) |
| *NKG* | Non Tillage |
| *SP* | Tillage |
| *SO* | Sampled Object |

**Figure S2**: field set up location Valthermond (peaty soils)

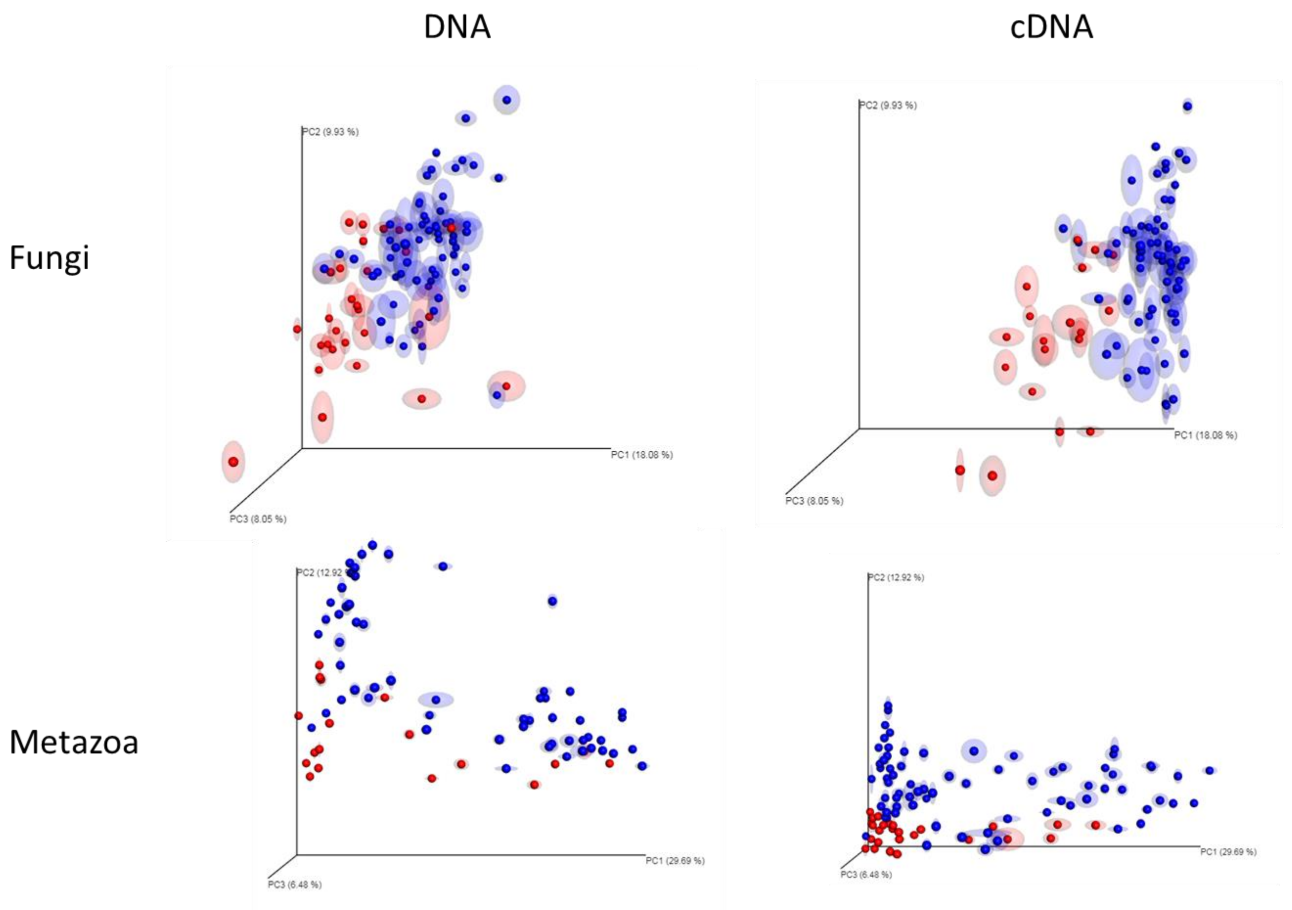


**Figure S3**: coordinate analysis (PCoA) plot with Bray-Curtis dissimilarity. Plots illustrating distances between communities in either DNA (n=104) A and C or cDNA (n=104) B and D. For Fungi A and B and Metazoa C and D. Distinguishing between Valthermond (Foley et al.) and Vredepeel (blue).

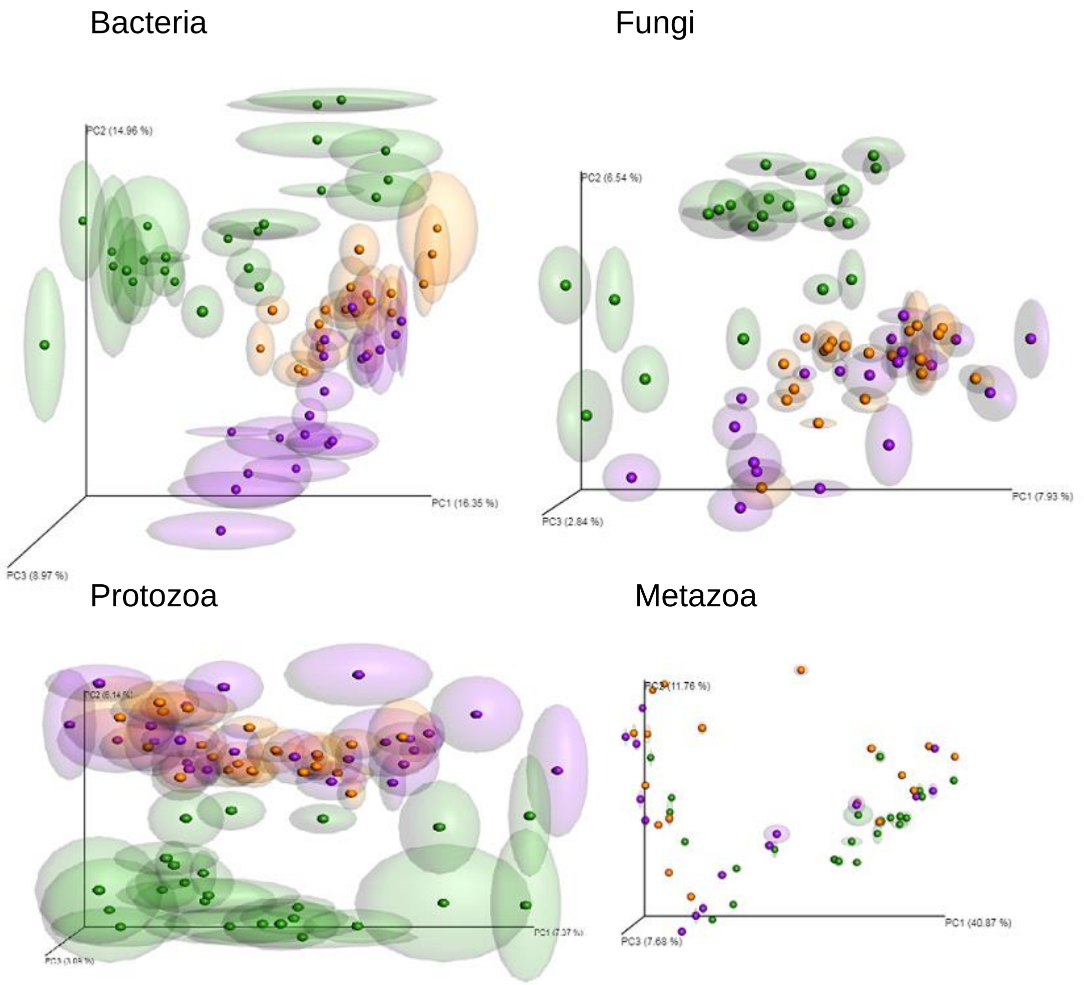


**Figure S4**: Principal coordinate analysis (PCoA) plot with Bray-Curtis dissimilarity. Plots illustrating distances between communities in all individual DNA samples from Vredepeel (n=72) for Bacteria, Fungi, Protozoa and Metazoa. Distinguishing between treatments: ConMin (purple) ConSlu (orange) and organic (green).

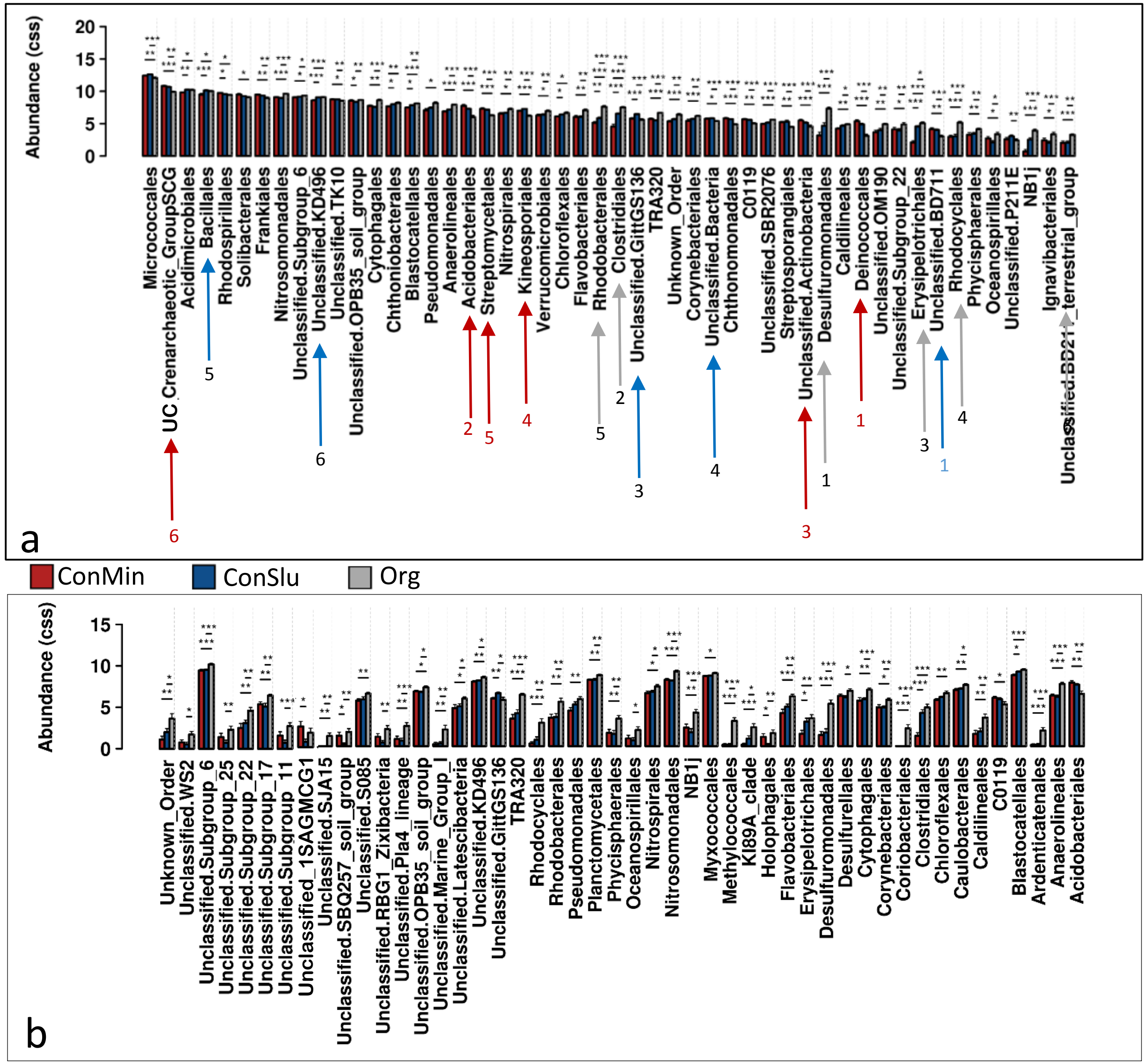


**Figure S5.1**: Differences in bacterial OTU abundance between the different soil management types (ConMin=red, ConSlu=blue and ORG=grey) for active (a) and total (b) bacterial communities. Asterisks represent statistically significant variance (ANOVA p-values; $*p < 0.05$, $**p < 0.01$, $***p < 0.001$) ANOVA. Arrows specify the order in which LefSe analysis categorises the indicative active orders.

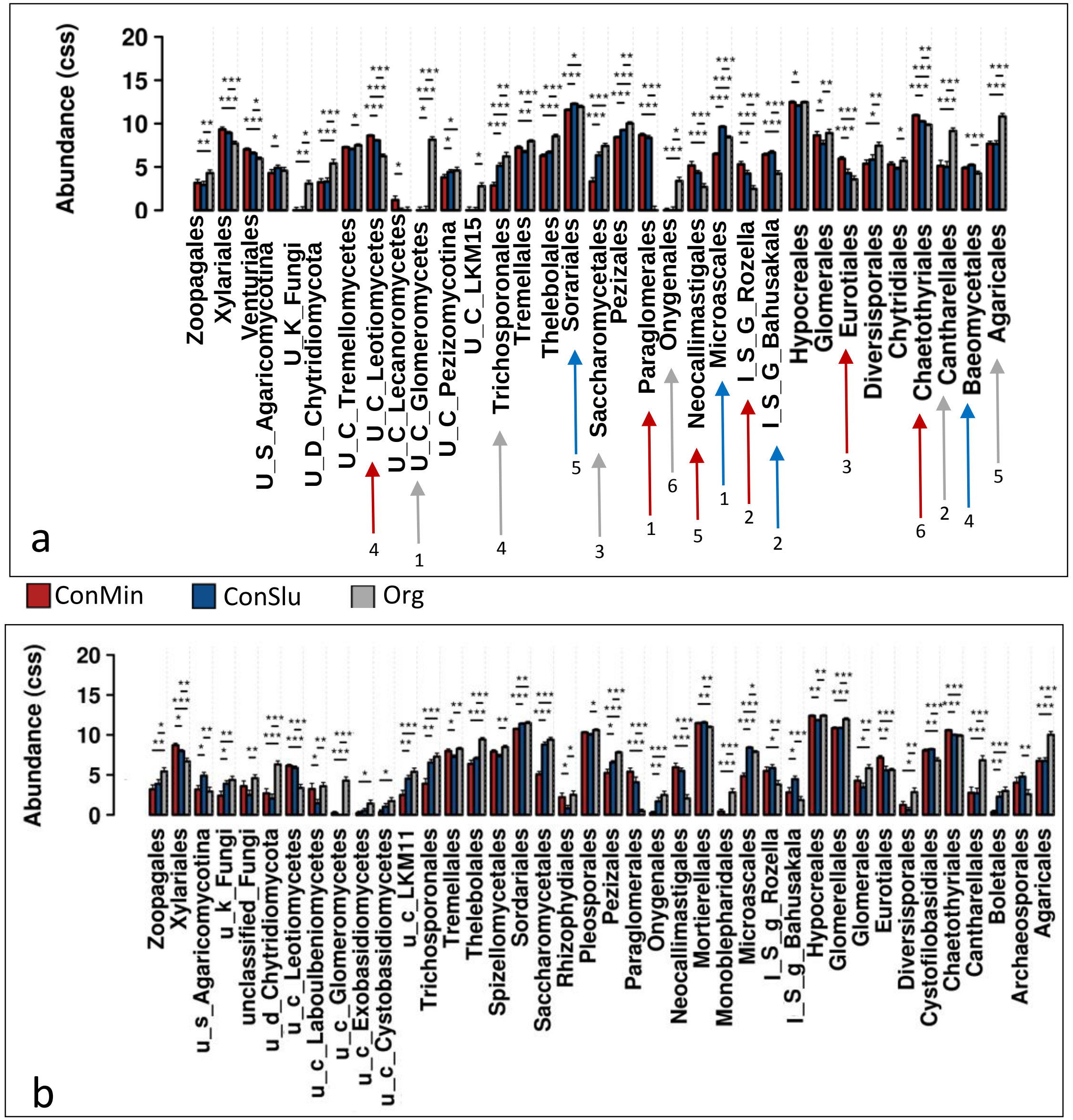


**Figure S5.2**: Differences in fungal OTU abundance between the different soil management types (ConMin=red, ConSlu=blue and ORG=grey) for active (a) and total (b) fungal communities. Asterisks represent statistically significant variance (ANOVA p-values; $*p < 0.05$, $**p < 0.01$, $***p < 0.001$) ANOVA. Arrows specify the order in which LefSe analysis categorises the indicative active orders.

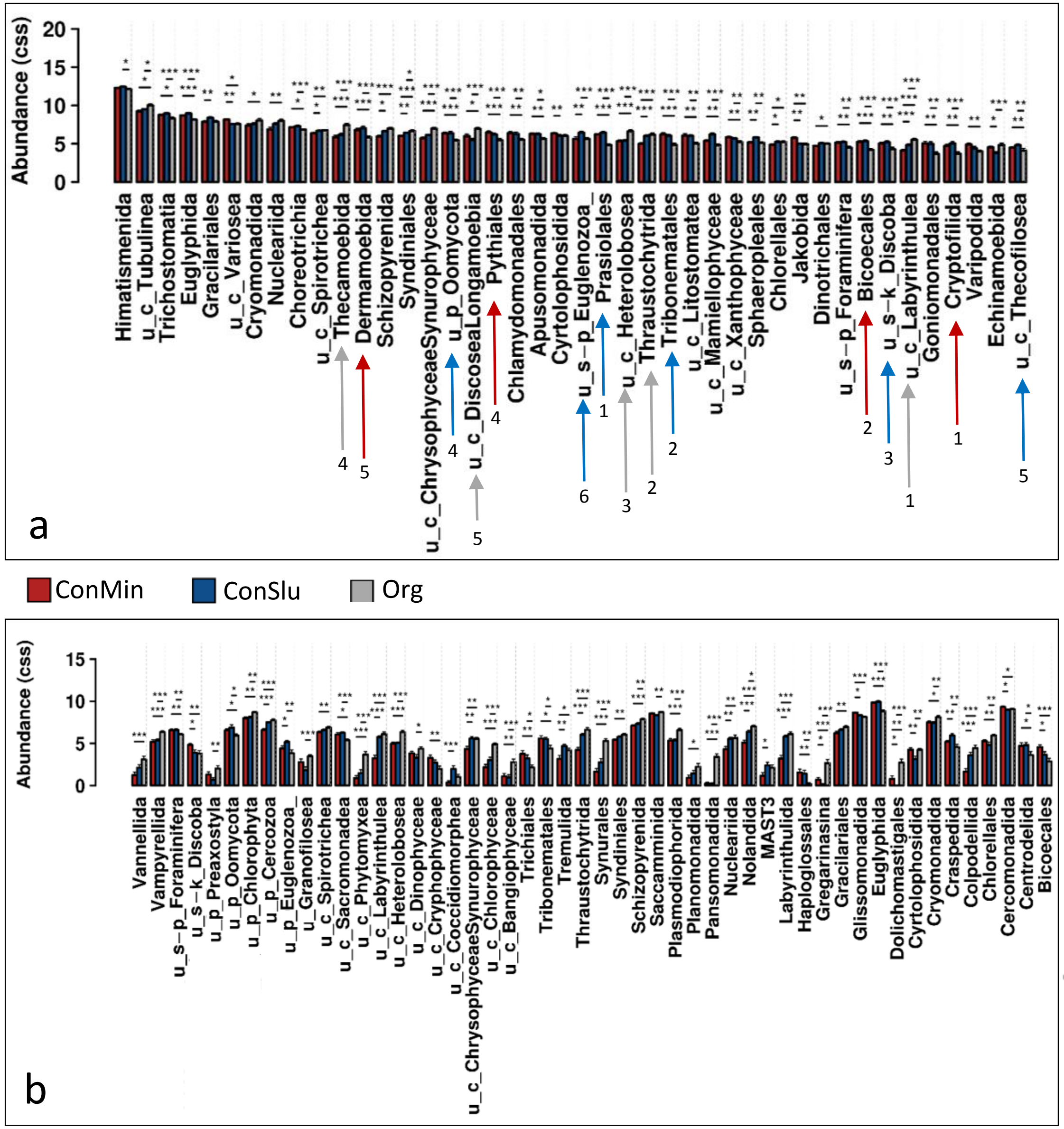


**Figure S5.3**: Differences in protist OTU abundance between the different soil management types (ConMin=red, ConSlu=blue and ORG=grey) for active (a) and total (b) protist communities. Asterisks represent statistically significant variance (ANOVA p-values; $*p < 0.05$, $**p < 0.01$, $***p < 0.001$) ANOVA. Arrows specify the order in which LefSe analysis categorises the indicative active orders.
